# Classification of the glyphosate target enzyme (5-enolpyruvylshikimate-3-phosphate synthase)

**DOI:** 10.1101/2020.05.27.118265

**Authors:** Lyydia Leino, Tuomas Tall, Marjo Helander, Irma Saloniemi, Kari Saikkonen, Suvi Ruuskanen, Pere Puigbò

## Abstract

Glyphosate is the most common broad-spectrum herbicide. It targets the key enzyme of the shikimate pathway, 5-enolpyruvylshikimate-3-phosphate synthase (EPSPS), which synthesizes three essential aromatic amino acids (phenylalanine, tyrosine and tryptophan) in plants. Because the shikimate pathway is also found in many prokaryotes and fungi, the widespread use of glyphosate may have unsuspected impacts on the diversity and composition of microbial communities, including the human gut microbiome. Here, we introduce the first bioinformatics method to assess the potential sensitivity of organisms to glyphosate based on the type of EPSPS enzyme. We have precomputed a dataset of EPSPS sequences from thousands of species that will be an invaluable resource to advancing the research field. This novel methodology can classify sequences from >90% of eukaryotes and >80% of prokaryotes. A conservative estimate from our results shows that 54% of species in the core human gut microbiome are sensitive to glyphosate.

## INTRODUCTION

Glyphosate is the most efficient and widely used nonselective herbicide. Historically, it was commercialized in the 1970s and then became the most inexpensive herbicide after the patent expired in 2000 ^1–3^. Since then, numerous generic glyphosate-containing herbicides have made glyphosate-based herbicides (GBHs) the most commonly used pesticides worldwide ^4^. The dominance of GBHs in the pesticide market is mainly attributed to the use of transgenic crops such as soy, corn and canola, of which nearly 90% are glyphosate-resistant varieties ^5^. In Europe, where transgenic crops are hardly grown, they are much used in no-till cropping, where weeds are eradicated by glyphosate prior to sowing. In addition, cereal, bean and seed crops are commonly desiccated by glyphosate before harvest.

The biochemical target enzyme for the herbicide glyphosate is 5-enolpyruvylshikimate-3-phosphate synthase (EPSPS) ^6^, the key enzyme of the shikimate pathway, which synthesizes the three essential aromatic amino acids (phenylalanine, tyrosine and tryptophan) in most prokaryotes, plants and fungi ^7,8^. Glyphosate is proclaimed safe for humans and other nontarget organisms because the shikimate metabolic pathway, inactivated by glyphosate, is not present in vertebrates. However, until recently, the presence of the shikimate pathway and diversity of EPSPS in many microbes have largely been ignored. As microbes are ubiquitous, associated with virtually all higher organisms, and essential in maintaining fundamental organismal functions ^9–11^, predicting the consequences of glyphosate use via its potential effects on the microbiome is challenging. The first step toward a more comprehensive understanding of how glyphosate affects higher organisms and biotic interactions involving microbes is to survey microbe susceptibilities to glyphosate.

The widespread use of glyphosate may have a strong impact on the diversity and community composition of both eukaryotes (mainly plants and fungi) and prokaryotes. Nearly all plant and most fungal populations are sensitive to glyphosate ^12,13^, but we are still only beginning to understand the effects of glyphosate on the gut microbiome. The first evidence has shown that glyphosate can affect the bee gut microbiota composition ^14^. An association between the use of glyphosate and antibiotic resistance has been suggested ^15,16^, though it is not clear in which direction, and thus, further studies are needed.

Here, we propose the first bioinformatic method to predict the glyphosate sensitivity/resistance of organisms based on the type of EPSPS enzyme. We have used this methodology to perform a comprehensive classification of the glyphosate target enzyme, EPSPS, based on sensitivity/resistance to glyphosate, taxonomic distribution and domain architectures. Our methodology classifies EPSPS enzymes into four different classes with differential sensitivities to glyphosate based on the presence and absence of amino acid markers in the active site ^17–24^. The classification of organisms based on the type of EPSPS enzyme will be extremely useful to assess species that are putatively sensitive or resistant to glyphosate. We have precomputed a dataset of EPSPS enzymes from thousands of species that will be an invaluable resource to test hypotheses in the field. This dataset includes 890 sequences from species in the core human gut microbiome, of which 54% are putatively sensitive to glyphosate (in a conservative estimate).

## RESULTS

### Classification of EPSPS enzymes based on sensitivity to glyphosate

EPSPS enzymes can be classified into four groups based on differential sensitivity to the herbicide glyphosate. This classification is based on the presence and absence of amino acid markers in the active site of the protein ^17–24^ (figure 1, supplementary figures 1–2). In general, species containing class I EPSPS sequences (alpha and beta) are sensitive to glyphosate, whereas species with class II sequences tend to be resistant ^19–22^ (supplementary table 1). EPSPS proteins belonging to classes III and IV putatively result in resistance to glyphosate ^23,24^ and are relatively rare in nature (<5% of the sequences), e.g., all class IV EPSPS sequences are found in one single actinobacteria clade (figure 2b and supplementary figure 3). In prokaryotes, the majority of species have class I enzymes and are thus sensitive to glyphosate (84% in archaea and 57% in bacteria), whereas class II enzymes (resistant to glyphosate) represent only 2% and 32% of archaeal and bacterial species, respectively. In eukaryotes, the majority of EPSPS proteins belong to class I, including 69% of viridiplantae species and 91% of fungi. Although a relatively large portion of eukaryotic species remain unclassified (especially in viridiplantae, 31%), the number of species potentially resistant to glyphosate based on amino acid markers is quite low (figure 2 and supplementary figures 4 and 5). As an example, we determined the EPSPS classes in symbiotic fungal endophytes living internally and asymptomatically within many grass species (figure 3). Recent studies have shown that glyphosate may alter this symbiotic relationship ^25^. EPSPS domains from endophytes were aligned with four reference sequences of EPSPS enzymes that contain the key amino acid markers of classes I-IV (supplementary tables 1–2). Although there are variations in the numbers and lengths of EPSPS-associated domains in endophytes, all EPSPS sequences indicated sensitivity to glyphosate (class I).

**Table 1.** List of common gut bacteria and their susceptibility to glyphosate in a taxonomic level of species. Species sensitive and resistant to glyphosate are highlighted in green and red respectively. The species that showed intraspecific variation are highlighted in blue and unclassified species are highlighted in black. If the EPSPS of a strain is not of any Class, the sensitivity is unknown. The total number of sequences is N=890.

| Sensitive to glyphosate (Class I, 100%) |  | Resistant to glyphosate (Class II , 100%) | Resistant to glyphosate (Class III, 100%) |
| --- | --- | --- | --- |
| *Actinomyces odontolyticus* (1) | *Parabacteroides merdae* (8) | *Anaerobaculum hydrogeniformans* (1) | *Alistipes putredinis* (3) |
| *Anaerofustis stercorihominis* (1) | *Bifidobacterium breve* (28) | *Blautia hansenii* (3) |  |
| *Anaerotruncus colihominis* (5) | *Bifidobacterium catenulatum* (8) | *Bryantella formatexigens* (1) | Some sensitive and resistant strains |
| *Bacteroides caccae* (5) | *Bifidobacterium dentium* (5) | *Clostridium asparagiforme* (2) | (Class I , 96% / Class II, 4%) |
| *Bacteroides capillosus* (1) | *Bifidobacterium longum* (102) | *Clostridium bartlettii* (2) | *Bifidobacterium adolescentis* (28): 27S + 1R |
| *Bacteroides cellulosilyticus* (4) | *Catenibacterium mitsuokai* (3) | *Clostridium scindens* (2) |  |
| *Bacteroides coprocola* (2) | *Citrobacter portucalensis* (5) | *Clostridium* sp. SS2/1 (2) | Some sensitive and unclassified strains |
| *Bacteroides coprophilus* (2) | *Clostridium methylpentosum* (1) | *Clostridium symbiosum* (1) | (Class I, 94% / No Class, 6%) |
| *Bacteroides eggerthii* (3) | *Clostridium nexile* (1) | *Coprococcus comes* (9) | *Fusobacterium nucleatum* (2): 1S +1U |
| *Bacteroides finegoldii* (4) | *Collinsella aerofaciens* (9) | *Coprococcus eutactus* (6) | *Bifidobacterium pseudocatenulatum* (46): 45S + 1U |
| *Bacteroides fragilis* (56) | *Desulfovibrio piger* (4) | *Dorea formicigenerans* (18) | *Clostridium leptum* (5): 4S + 1U |
| *Bacteroides intestinalis* (11) | *Enterobacter cancerogenus* (1) | *Dorea longicatena* (11) |  |
| *Bacteroides ovatus* (14) | *Escherichia coli* (112) | *Enterococcus casseliflavus* (2) | Some resistant and unclassified strains |
| *Bacteroides plebeius* (11) | *Eubacterium siraeum* (4) | *Eubacterium hallii/Anaerobutyrum hallii* (9) | (Class II, 79% / No Class, 21%) |
| *Bacteroides* sp. 2_2_4 (1) | *Faecalibacterium prausnitzii* (53) | *Helicobacter canadensis* (1) | *Clostridium* sp. L2/50 (2): 1R + 1U |
| *Bacteroides* sp. 3_2_5 (1) | *Fusobacterium mortiferum* (1) | *Helicobacter cinaedi* (2) | *Enterococcus faecalis* (4): 3R + 1U |
| *Bacteroides* sp. 4_3_47FAA (1) | *Fusobacterium ulcerans* (1) | *Helicobacter pullorum* (1) | *Ruminococcus torques* (8): 7R + 1U |
| *Bacteroides* sp. 9_1_42FAA (2) | *Parabacteroides* sp. 2_1_7 (1) | *Helicobacter winghamensis* (1) |  |
| *Bacteroides* sp. D1 (1) | *Prevotella copri* (27) | *Lactobacillus brevis subsp. gravesensis* (1) | Unclassified strains (No Class, 100%) |
| *Bacteroides thetaiotaomicron* (13) | *Providencia alcalifaciens* (1) | *Lactobacillus buchneri* (1) | *Bacteroides pectinophilus* (1) |
| *Bacteroides uniformis* (10) | *Providencia rustigianii* (1) | *Lactobacillus hilgardii* (1) | *Butyrivibrio crossotus* (2) |
| *Bacteroides vulgatus* (6) | *Providencia stuartii* (1) | *Lactobacillus plantarum* (2) | *Clostridium spiroforme* (1) |
| *Bifidobacterium bifidum* (22) | *Ruminococcus bromii* (14) | *Lactobacillus ruminis* (2) | *Eubacterium rectale* (26) |
| *Fusobacterium varium* (1) | *Subdoligranulum variabile* (3) | *Listeria grayi* (1) | *Eubacterium ventriosum* (7) |
| *Holdemania filiformis* (1) |  | *Mitsuokella multacida* (3) | *Fusobacterium necrophorum* (1) |
| *Mollicutes bacterium* (3) |  | *Ruminococcus gnavus* (17) | *Fusobacterium periodonticum* (1) |
| *Parabacteroides distasonis* (10) |  | *Ruminococcus lactaris* (3) | *Proteus penneri* (1) |
| *Parabacteroides johnsonii* (4) |  | *Ruminococcus obeum* (18) | *Providencia rettgeri* (1) |
|  |  |  | *Roseburia intestinalis* (9) |

(n): Number of sequences; R: glyphosate resistant sequences; S: glyphosate sensitive sequences, U: unclassified.

**Figure 1.**
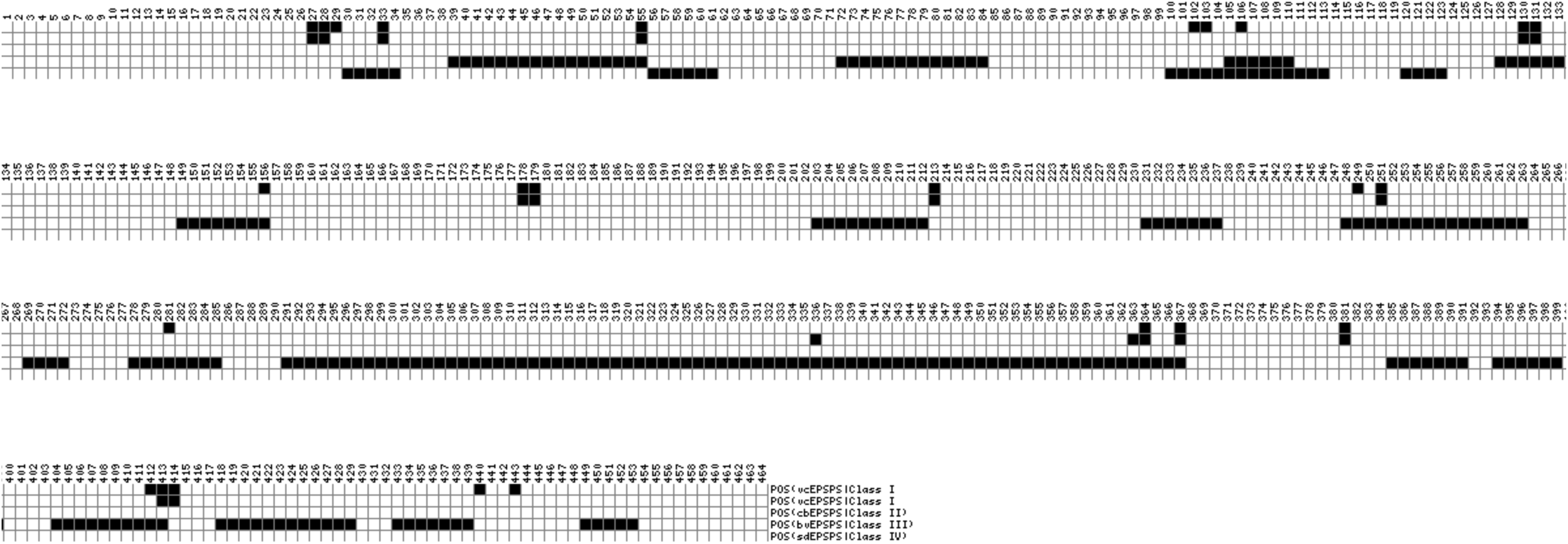
Position of amino acid markers in the reference EPSPS sequences. Amino acid markers of EPSPS classes I (EPSPS from *Vibrio cholerae*; vcEPSPS), II (EPSPS from *Coxiella burnetii*; cbEPSPS), III (EPSPS from *Brevundimonas vesicularis*; bvEPSPS) and IV (EPSPS *Streptomyces davawensis*; sdEPSPS) are shown in black. The complete list of amino acids is found in supplementary table 1 and supplementary figure 2. This figure was made with the web server https://matrix2png.msl.ubc.ca/.

**Figure 2.**
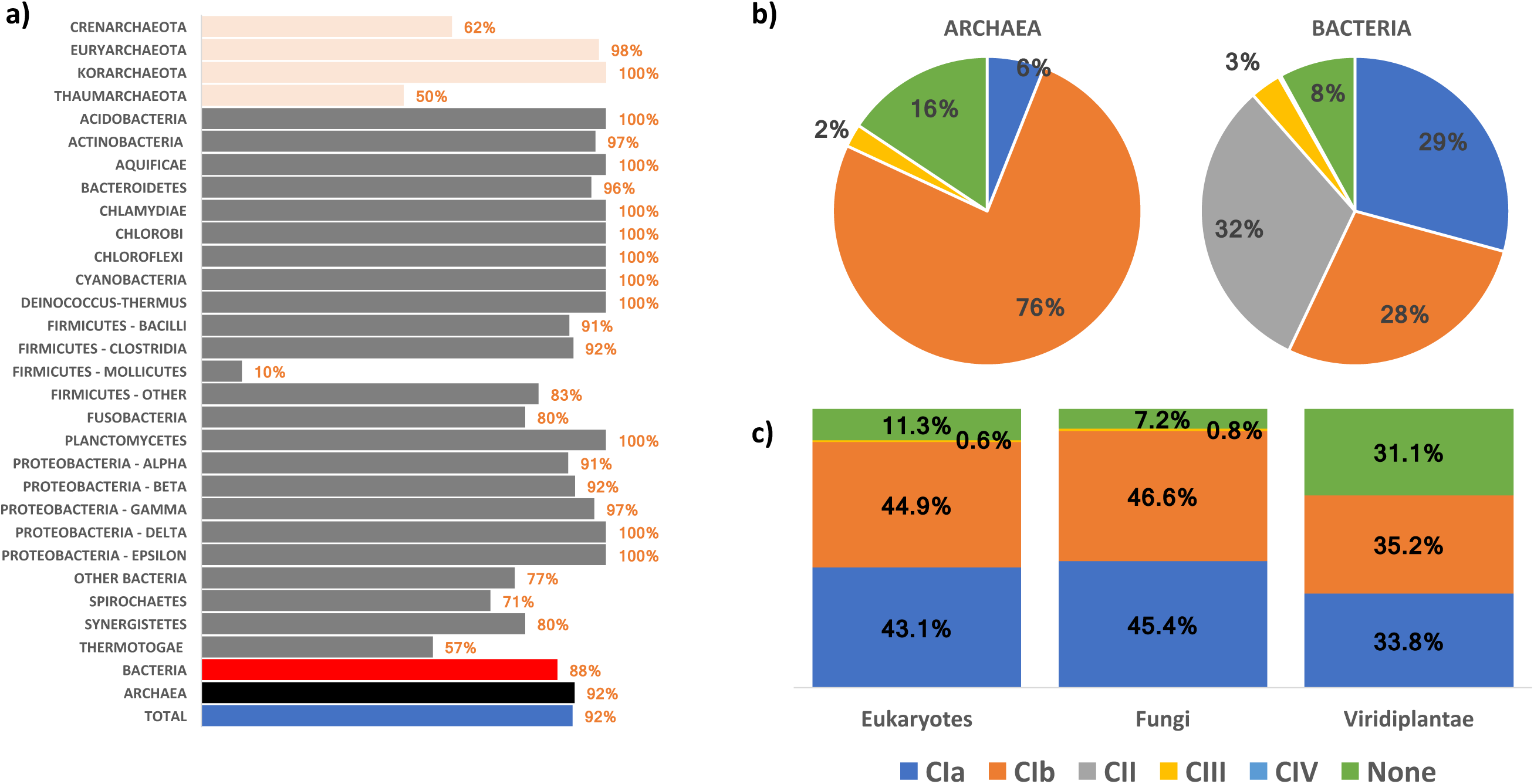
Taxonomic distribution and classification of EPSPS proteins. a) Percentage of the EPSPS proteins in prokaryotic sequences within each taxon. EPSPS is present in 652 out of the 678 genomes analyzed. The data were obtained from the COG database. EPSPS proteins belong to COG0128 (category E; amino acid transport and metabolism). b) Percentages of classes I, II, III, and IV and unclassified (none) in prokaryotes (source of sequences: COG database). c) Percentages of classes I, II, III, and IV and unclassified (none) in eukaryotes, fungi and viridiplantae (source of sequences: UniProt database).

**Figure 3.**
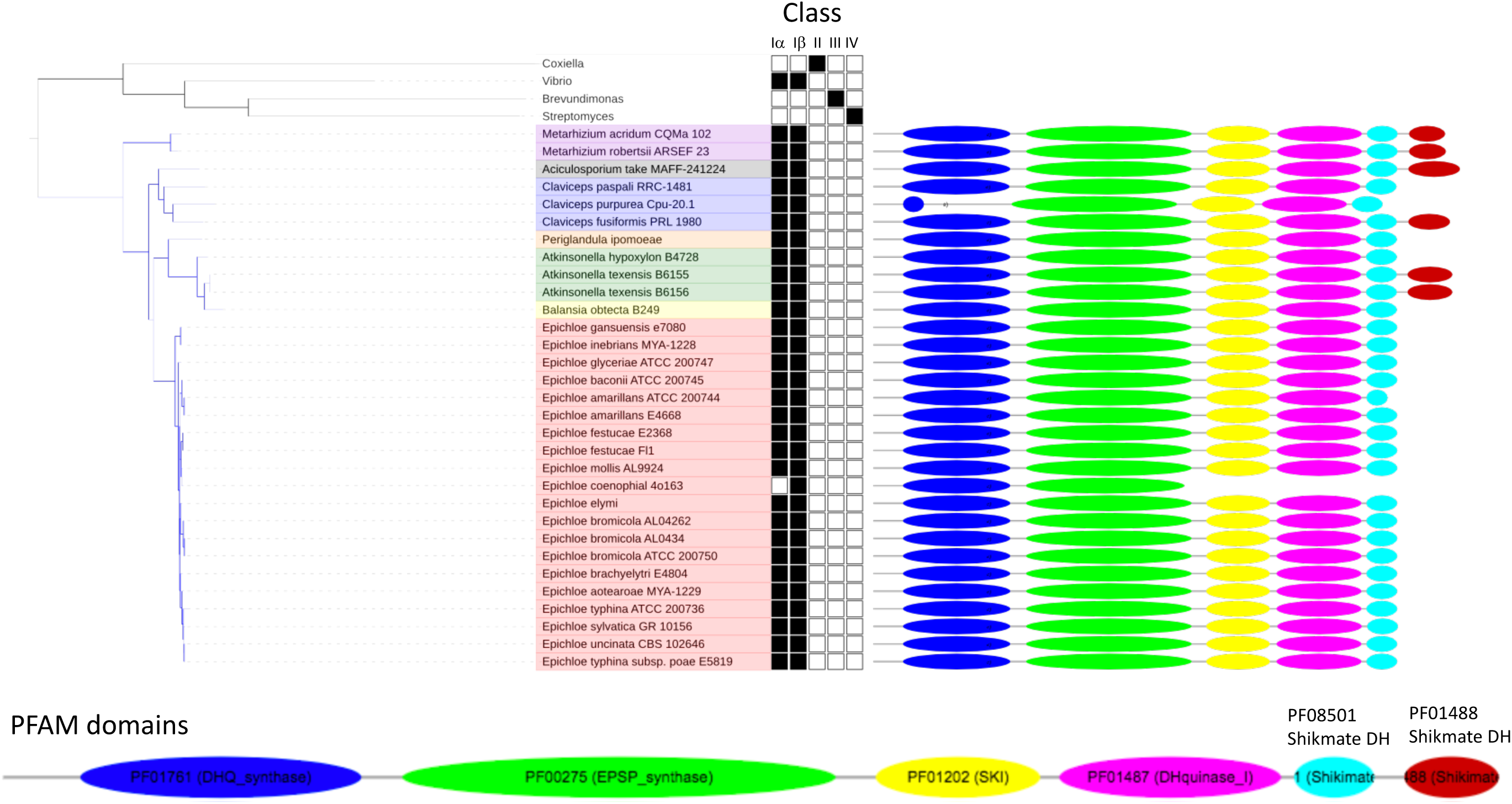
Identification of the EPSPS class in endophytes. The EPSPS enzymes of endophytes are mapped onto the reference sequences (class I, vcEPSPS; class II, cbEPSPS; class III, bvEPSPS; and class IV, sdEPSPS) to determine the potential sensitivity to glyphosate. Endophyte EPSPS sequences were obtained from http://www.endophyte.uky.edu.

**Figure 4.**
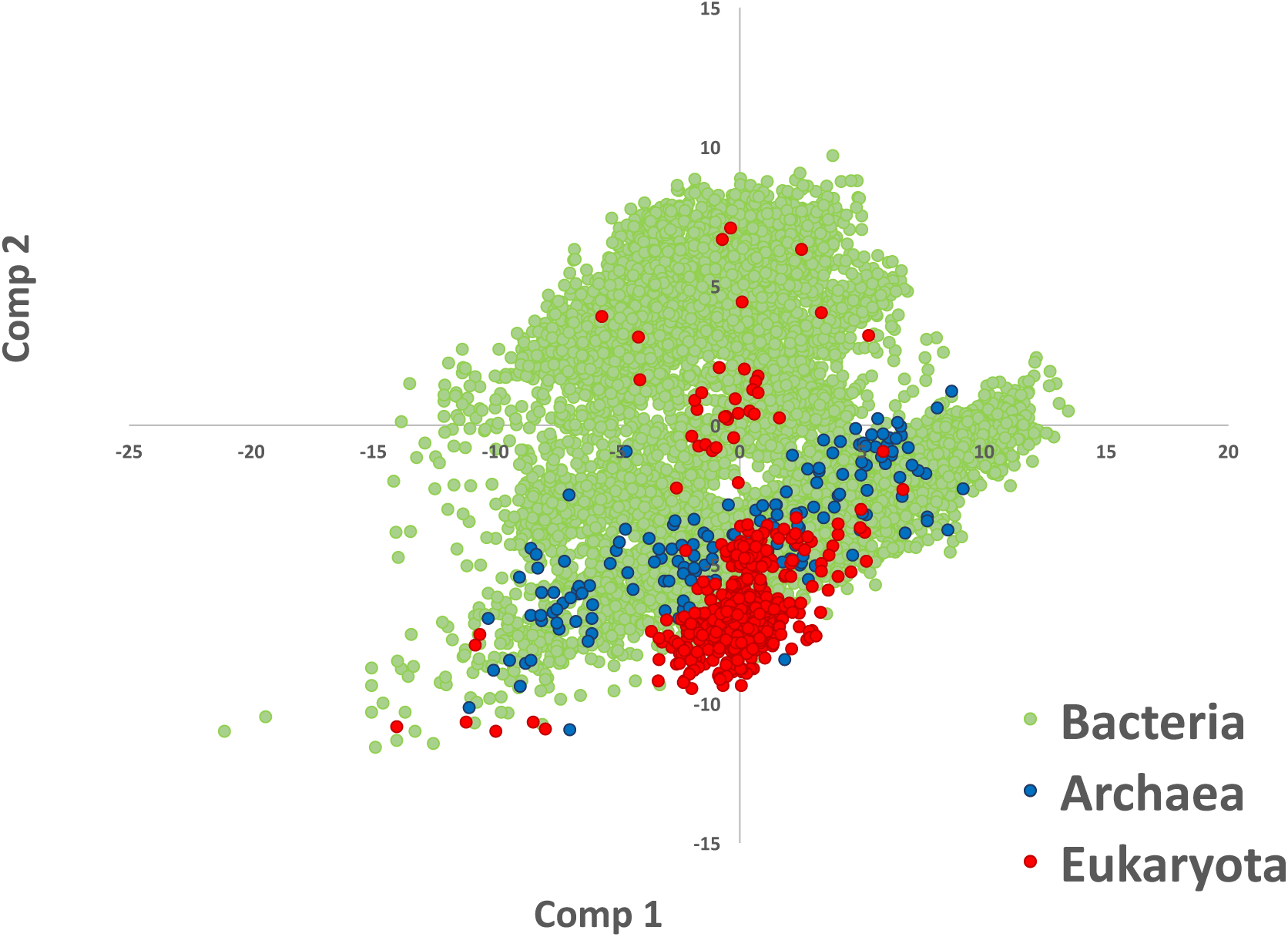
Principal components analysis (PCA) plot of dipeptides. The plot shows the first and second components of the PCA from ∼10,000 EPSPS proteins of archaea (blue), bacteria (green) and eukaryotes (red). Additional clusters from this plot, based on taxonomy and EPSPS class, are available in supplementary figures 3–4.

**Figure 5.**
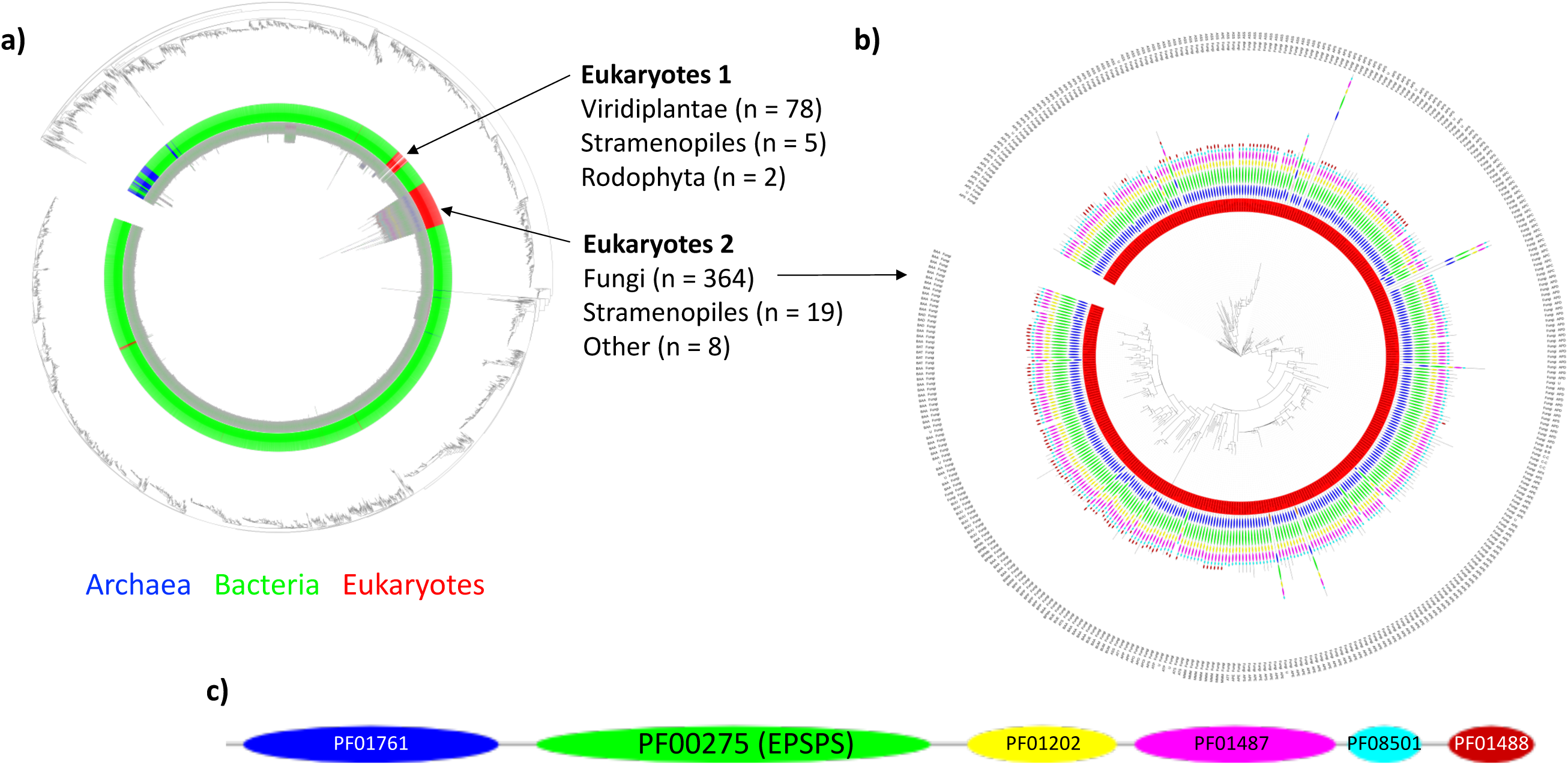
Phylogenetic tree of the EPSPS domain made from ∼10,000 sequences (>350 AA) obtained from the Uniprot database. a) Phylogenetic tree including archaea (blue), bacteria (green) and eukaryotes (red) b) Phylogenetic tree of the fungal clade shows the conservation of the multidomain structure of the EPSPS protein. c) Example of a common multidomain architecture in a fungal EPSPS protein that includes the domains EPSPS (PF00275) and its associated domains PF01488 (Shikimate_DH), PF08501 (Shikimate_DH_N), PF01487 (DH_Quinase_I), PF01202 (SKI) and PF01761 (DHQS). The majority of bacteria and plants contain only the EPSPS domain (PF00275).

A large portion of EPSPS proteins (especially in prokaryotes) do not belong to any of the four known classes and are thus hereinafter termed unclassified EPSPS (figure 4, supplementary tables 3–4). Moreover, an analysis of dipeptides of EPSPS domains shows taxonomic and class differences among sequences (figure 4, supplementary figures 4 and 5). Both eukaryotic and archaeal species are merged within the larger cluster of bacterial species, but it is possible to differentiate two independent clusters for fungi and another two clusters for viridiplantae. This pattern is also shown in certain bacterial groups, e.g., in Proteobacteria, Firmicutes, Actinobacteria and cyanobacteria. Further empirical studies are needed to identify novel amino acid markers that determine the potential sensitivity of unclassified EPSPS proteins to glyphosate. Therefore, our dataset of more than 50,000 EPSPS proteins obtained from public databases ^26,27^ and classified according to the potential sensitivity to the herbicide glyphosate (classes I–IV or unclassified) will be highly useful for future studies. This dataset is open and freely available at https://ppuigbo.me/programs/EPSPSClass. Moreover, a web server is also offered to compute the EPSPS class of a protein from its raw amino acid sequence, which will be highly useful in identifying new classes of EPSPS proteins.

### Taxonomic distribution of the EPSPS enzymes

EPSPS enzymes are present in three domains of life: archaea, bacteria and eukaryotes (mostly in plants and fungi) ^7,8^. A phylogenetic analysis of the EPSPS domain (>10,000 sequences) shows that none of bacteria, archaea or eukaryotes form a monophyletic group (Figure 5a and supplementary tables 5–6). In eukaryotes, EPSPS is present in fungi, viridiplantae, stramenopiles and rhodophyta in two independent paraphyletic clusters. The first cluster of eukaryotes (Eukaryotes 1) contains single-domain sequences corresponding to viridiplantae (main taxa), stramenopiles and rhodophyta, whereas the second cluster (Eukaryotes 2) contains multidomain proteins (including the EPSPS domain and associated domains) from fungi and a few stramenopiles (figure 5bc). The phylogenetic analysis is based on alignment of only the EPSPS domain (see the Materials and Methods); thus, the independent eukaryotic clades may not be explained by different domain architectures. Moreover, there are additional scattered eukaryotic sequences within bacterial groups that may be the product of contamination during genome sequencing (e.g., a simple BLAST analysis showed that the EPSPS of the winter moth is the product of bacterial contamination –data not shown) and putative ancient horizontal gene transfers (e.g., a relatively large cluster including several rosids) from bacterial species (supplementary figure 6). An analysis of COG0128 from the Clusters of Orthologous Groups (COG) database ^28^, which contains precomputed orthologous genes of more than 700 prokaryotes ^27^, shows that 92% of archaea and 88% of bacteria have at least one copy of the EPSPS protein (figure 5a). Nevertheless, the enzyme is remarkably infrequent in some bacterial groups, such as Mollicutes (present in only 10% of species) and Thermotoga (present in only 57% of species).

### EPSPS domain architectures and analysis of the EPSPS-associated domains

EPSPS proteins include (by definition) the EPSPS domain, but in several species, the EPSPS protein contains additional associated domains (e.g., fungal species) (figure 5c and supplementary table 5). We analyzed ∼10,000 protein sequences from the Pfam database ^26^ that contain the domain PF00275 (EPSPS). The EPSPS domain is ∼450 amino acids long and is present in 62 domain architectures and as a single domain in most prokaryotes and viridiplantae (Pfam data ^26^). In fungi, the EPSPS domain forms part of multidomain proteins (usually more than 5 domains) that are ∼1,300 amino acids long (figure 5c and supplementary table 6). Moreover, in a small number of bacterial and viridiplantae species, the EPSPS protein contains the EPSPS domain and an additional associated domain. The most common EPSPS-associated domains are shikimate pathway (SKI, DHQ synthase, DHquinase I) and aromatic amino acid synthesis (Shikimate dh N, PDH) enzymes and promiscuous domains (HTH 3). Additional EPSPS-associated domains exhibit functions such as DNA modification, gene expression and other enzymatic activities (supplementary table 6). Overall, the EPSPS-associated domains can be classified into shikimate (shikimate pathway domains), enzymatic (domains with catalytic function), expression (domains whose products are needed in controlling gene expression) and structural functions (domains that do not have catalytic function, e.g., binding sites, histones and helix-turn-helix domains).

### Survey of the resistance and sensitivity to glyphosate in the human gut microbiomes

To test this new resource in a real-world scenario, we analyzed EPSPS sequences of 890 strains from 101 common bacterial species in the human gut microbiome ^29^. Our results suggest that EPSPS sensitivity to glyphosate is quite conserved within bacterial species in the human gut microbiome (supplementary table 7). Fifty-four percent of species have an EPSPS in class I, i.e., potentially sensitive to glyphosate (table 1). The core gut microbiome contains 75 species together representing 22-47% of the total species abundance in the gut microbiome, which contains approximately 160 varying species^29^. Therefore, with a conservative calculation, 12-26% of bacterial species in the human microbiome might be sensitive and thus affected by glyphosate. In addition, 29% of species in the core microbiome are likely to be resistant to glyphosate (class II or III), and 17% of the species are still unclassified or contain some unclassified strains (table 1). Class IV EPSPS enzymes are not found in the microbiome dataset ^29^, as they are almost exclusive to streptomyces and a few other actinobacteria (table 3).

Within the ten most frequent bacterial species in the core human gut microbiome ^29^, four species are resistant to glyphosate (*Dorea formicigenerans; Clostridium* sp. SS2-1; *Eubacterium hallii; Coprococcus comes* SL7 1), four are sensitive (*Faecalibacterium prausnitzii SL3-3; Bacteroides vulgatus* ATCC 8482; *Bacteroides uniformis* and *Bacteroides* sp. *9 1* 42FAA), and two are unclassified (*Roseburia intestinalis M50 1* and *Eubacterium rectale* M104 1). The list of genera with strains sensitive to glyphosate includes (all sequences from the first 18 genera are sensitive to glyphosate) Actinomyces, Anaerofustis, Anaerotruncus, Catenibacterium, Citrobacter, Collinsella, Desulfovibrio, Enterobacter, Faecalibacterium, Holdemania, Methanobrevibacter, Mollicutes, Parabacteroides, Prevotella, Subdoligranulum, Escherichia, Actinomyces, Anaerofustis, Bacteroides, Bifidobacterium, Clostridium, Eubacterium, Ruminococcus, Providencia, and Fusobacterium. Moreover, in some genera, the sensitivity to glyphosate may vary widely, e.g., all EPSPS sequences from *Ruminococcus bromii* strains (n=14) belong to class I, whereas those from *Ruminococcus gnavus* strains (n=17) belong to class II. We also detected intraspecific variation in EPSPS sequences in 7% of all studied species (table 1).

## DISCUSSION

A large proportion of bacteria in the gut microbiome ^29^ are susceptible to glyphosate (class I); thus, the intake of glyphosate may severely affect the composition of the human gut microbiome. Our analysis suggests that the proportion of species susceptible to glyphosate is at least 12-26% of the total species in the human gut. Thus, the use of glyphosate may provide a competitive advantage to bacteria resistant to glyphosate over sensitive bacteria. Although data on glyphosate residues in human gut systems are still lacking ^29^, our results suggest that glyphosate residues decrease bacterial diversity and modulate bacterial species composition in the gut. Nevertheless, other studies have shown the impact of glyphosate on microbiomes ^14,30,31^. We may assume that long-term exposure to glyphosate residues leads to the dominance of resistant strains in the bacterial community. Moreover, some sensitive strains may become resistant to glyphosate through accumulation of mutations in the EPSPS domain or acquisition of a resistance gene *via* horizontal gene transfer. In plants (*Chloris virgata*) and bacteria (*Escherichia coli* and *Salmonella typhimurium*), changing Pro106 to Ser reduces the organism’s sensitivity to glyphosate ^32,33^. Although Pro106 does not molecularly directly interact with glyphosate, its substitution with different amino acids results in structural changes in the active site, inhibiting glyphosate action ^34^. In plants, Gly101, Thr102, Pro106, Gly144 and Ala192 mutations can provide diverse levels of resistance ^34^. In *E. coli* and comparable bacteria, the corresponding positions in EPSPS (supplementary figure 7) are Gly96, Thr97, Pro101, Gly137 and Ala183.

In vitro studies have revealed that species such as *Bacteroides vulgatus, Bifidobacterium adolescentis, Enterococcus faecalis, Staphylococcus aureus* and *Lactobacillus buchneri* treated with Roundup UltraMax® were moderate to highly sensitive to glyphosate (minimal inhibitory concentration (MIC) = 0.075 – 0.600 mg/ml), whereas *Escherichia coli* and *Salmonella typhimurium* were relatively unreactive (MIC values = 1.200 and 5.000 mg/ml) ^31^. Our results indicate that *B. vulgatus, B. adolescentis, E. coli* and *S. typhimurium* are mostly sensitive to glyphosate according to their EPSPS (table 1). Results from Shehata et al. do not disprove this, even though the last two species did show high tolerance in vitro ^*31*^. However, *E. faecalis, L. buchneri* and *S. aureus*, which we predict to be resistant according to the amino acid markers in EPSPS, are actually fairly sensitive to glyphosate in vitro ^*31*^. This suggests that some other factors can modulate the sensitivity of species to glyphosate-related products. These can include surfactants in different products (the use of Roundup UltraMax® could have toxic effects on glyphosate-resistant bacteria)

TThe complexity of glyphosate’s effects on the gut microbiota not only is dependent on the direct impacts of glyphosate blocking (or not) the EPSPS enzyme but also has an indirect effect on bacterial interactions. Numerous bacteria express antagonistic activity *via* bacteriocins ^31,35^. The impact of glyphosate on antagonist species is beyond the scope of this article, but it may potentially disrupt microbial biofilms ^36^. Projecting the exact effects of glyphosate on individual microbes or interactions among them in complex communities and the potential cascading effects on higher trophic levels is not straightforward. Thus, the bioinformatics tool and the precomputed dataset presented in this article will be highly useful to elucidate the impact of glyphosate on human health. To determine the actual impact of glyphosate on the human gut microbiota and other organisms, further empirical studies are needed (1) to reveal glyphosate residues in food, (2) to determine the effects of pure glyphosate and commercial formulations on microbiomes and (3) to assess the extent to which our EPSPS amino acid markers predict bacterial susceptibility to glyphosate in in vitro and real-world scenarios.

## CONCLUSIONS

We have introduced a comprehensive classification of EPSPS proteins based on their potential sensitivity to the broad-spectrum herbicide glyphosate. This classification of organisms based on the type of EPSPS enzyme can be utilized to test several hypotheses related to the use of glyphosate and related commercial products. Moreover, this novel resource that includes a precomputed dataset of more than 50,000 proteins and a web server to determine EPSPS protein classes will be highly useful in further research. A conservative estimate from our results shows that 54% of species in the core of the human gut microbiome are sensitive to glyphosate, which represents approximately a quarter of the total number of bacterial species in the gut.

## MATERIAL AND METHODS

### Protein sequences and domain architectures

We obtained 10,231 protein sequences annotated 5-enolpyruvylshikimate-3-phosphate synthase (EPSPS) enzymes from the Pfam database (PF00275) ^37^. Domain architectures and taxonomical annotations were obtained from the Pfam ^37^ and Uniprot ^38^ databases, respectively. We also obtained 37 protein sequences, putatively resistant to glyphosate, from the International Survey of Herbicide-Resistant Weeds ^39^ and 679 protein sequences of prokaryotes from the database of Clusters of Orthologous Groups (COG0128) ^27^.

### Alignment and phylogenetic tree construction

A phylogenetic tree of the EPSPS synthase was built from protein sequences containing at least 350 amino acids to filter out severely truncated sequences. We aligned the sequences with the program muscle (-maxiters 2) ^40^ and refined the alignment with the program gblocks ^41^ with the minimal length of a block set at 6 amino acid positions, and the maximum number of allowed contiguous nonconserved amino acid positions was set at 20. The final alignment contained 152 positions from 6 selected blocks. This alignment was used to reconstruct a phylogenetic tree with the program fasttree ^42^ with default parameters.

### Principal components analysis of dipeptides

We obtained the frequencies of pairs of consecutive amino acids (dipeptides) of each protein sequence and performed a principal components analysis (PCA), using the prcomp function of the r software package ^43^. The resulting PCA plot was analyzed based on taxonomy and EPSPS classification.

### Classification of the EPSPS

EPSPS enzymes can be classified into classes I (alpha or beta) ^19,20^, class II ^21,22^, III ^23^ and IV ^24^ based on the presence of amino acid markers (classes I, II and IV) and motifs (class III). Markers and motifs used to classify EPSPS enzymes into one of the four classes are based on the amino acid positions of EPSPS enzymes from *Vibrio cholerae* (vcEPSPS, class I), *Coxiella burnetii* (cbEPSPS, class II), *Brevundimonas vesicularis* (bvEPSPS, class III), and *Streptomyces davawensis* (sdEPSPS, class IV) (supplementary table 5 and 6). To classify an EPSPS enzyme, we performed pairwise alignments of the query sequence and each reference sequence (vcEPSPS, cbEPSPS, bvEPSPS and sdEPSPS). An enzyme is classified as class I, class II and/or class IV if it contains all the amino acid markers from the respective reference sequence(s) ^19^ and classified as class III if it contains at least one complete motif (out of 18) from the sdEPSPS sequence ^20,24^.

### Web server and datasets

A web server to determine the EPSPS class is open and freely available at http://ppuigbo.me/programs/EPSPSClass. The *EPSPSClass* web server requires only a query protein sequence in FASTA format to determine the putative EPSPS class (I, II, III or IV) and to calculate an identity percentage for each class. However, the server is not limited to these classes, and users can easily test their own reference sequence and amino acid markers. Moreover, a series of protein datasets have been automatically precomputed from different databases, such as the UniProt ^38^, COG^27^, PDB ^44^ and NCBI ^45^ databases. These datasets of predictions of resistance and sensitivity to glyphosate for more than 50,000 EPSPS proteins are available from the server home page.

### Search for EPSPS sequences in the human gut microbiome

The target human gut bacterial species and strains were chosen according to supplementary tables 5, 8 and 12 provided by Qin et al. 2010 ^29^. We obtained a list of 75 nonredundant human gut bacterial species with >1% genome coverage by Illumina reads in >50% of the cohort individuals (n=124) ^29^ and >10% genome coverage in any number of individuals. Moreover, we added EPSPS sequences from additional strains to the dataset to determine the intraspecific diversity in EPSPS. We searched for additional human gut bacteria EPSPS sequences through protein BLAST searches ^46^ using vcEPSPS (supplementary table 2) as a reference. Out of the thousands of putative EPSPS sequences from the BLAST results, we selected only putative complete EPSPS sequences from bacterial strains (i.e., multispecies or partial sequences were excluded) belonging to the gut microbiome. The resulting sequences were analyzed with our *EPSPSClass* web server, which compares a query sequence against reference sequences for classes I, II, III and IV (supplementary table 2), to determine an identity percentage to each EPSPS class. Our final dataset (available from the server main page) contains the EPSPS classification for 890 strains from 101 species of prokaryotes.

## ACKNOWLEDGMENTS

This research is supported by funds from the Turku Collegium for Science and Medicine (PP, TT) and by the Academy of Finland (MH, grant #311077).

