## Supplementary Material for "Classification of the glyphosate target enzyme (5-enolpyruvylshikimate-3-phosphate synthase)"

### SUPPLEMENTARY TABLES

**Supplementary table 1. Alignment of EPSPS reference sequences.** Conserved amino acids are highlighted in blue (Class Iα), red (Class Iβ), green (Class II), yellow (Class III) and grey (Class IV).

| Positions | Class Iα Ref. vcEPSPS | Class Iβ Ref. vcEPSPS | Class II Ref. cbEPSPS | Class III Ref. bvEPSPS | Class IV Ref. sdEPSPS |
|---|---|---|---|---|---|
| **1** | -,- | -,- | -,- | 1,M | -,- |
| **2** | -,- | -,- | -,- | 2,M | -,- |
| **3** | -,- | -,- | -,- | 3,M | -,- |
| **4** | -,- | -,- | -,- | 4,G | -,- |
| **5** | -,- | -,- | -,- | 5,R | -,- |
| **6** | 1,M | 1,M | -,- | 6,A | -,- |
| **7** | 2,E | 2,E | 1,M | 7,K | -,- |
| **8** | 3,S | 3,S | 2,D | 8,L | -,- |
| **9** | 4,L | 4,L | 3,Y | 9,T | -,- |
| **10** | 5,T | 5,T | 4,Q | 10,I | -,- |
| **11** | 6,L | 6,L | 5,T | 11,I | -,- |
| **12** | 7,Q | 7,Q | 6,I | 12,P | 1,M |
| **13** | 8,P | 8,P | 7,P | 13,P | 2,P |
| **14** | 9,I | 9,I | 8,S | 14,G | 3,V |
| **15** | 10,E | 10,E | 9,Q | 15,K | 4,A |
| **16** | 11,L | 11,L | 10,G | 16,P | 5,D |
| **17** | 12,I | 12,I | 11,L | 17,L | 6,I |
| **18** | 13,S | 13,S | 12,S | 18,T | -,- |
| **19** | 14,G | 14,G | 13,G | 19,G | -,- |
| **20** | 15,E | 15,E | 14,E | 20,R | -,- |
| **21** | 16,V | 16,V | 15,I | 21,A | -,- |
| **22** | 17,N | 17,N | 16,C | 22,M | -,- |
| **23** | 18,L | 18,L | 17,V | 23,P | -,- |

| | | | | | |
|---|---|---|---|---|---|
| **24** | 19,P | 19,P | 18,P | 24,P | 7,P |
| **25** | 20,G | 20,G | 19,G | 25,G | 8,G |
| **26** | 21,S | 21,S | 20,D | 26,S | 9,S |
| **27** | 22,K | 22,K | 21,K | 27,K | 10,K |
| **28** | 23,S | 23,S | 22,S | 28,S | 11,S |
| **29** | 24,V | 24,V | 23,I | 29,I | 12,I |
| **30** | 25,S | 25,S | 24,S | 30,T | 13,T |
| **31** | 26,N | 26,N | 25,H | 31,N | 14,A |
| **32** | 27,R | 27,R | 26,R | 32,R | 15,R |
| **33** | 28,A | 28,A | 27,A | 33,A | 16,A |
| **34** | 29,L | 29,L | 28,V | 34,L | 17,L |
| **35** | 30,L | 30,L | 29,L | 35,L | 18,F |
| **36** | 31,L | 31,L | 30,L | 36,L | 19,L |
| **37** | 32,A | 32,A | 31,A | 37,A | 20,A |
| **38** | 33,A | 33,A | 32,A | 38,G | 21,A |
| **39** | 34,L | 34,L | 33,I | 39,L | 22,A |
| **40** | 35,A | 35,A | 34,A | 40,A | 23,A |
| **41** | 36,S | 36,S | 35,E | 41,K | 24,D |
| **42** | 37,G | 37,G | 36,G | 42,G | 25,G |
| **43** | 38,T | 38,T | 37,Q | 43,T | 26,V |
| **44** | 39,T | 39,T | 38,T | 44,S | 27,T |
| **45** | 40,R | 40,R | 39,Q | 45,R | 28,T |
| **46** | 41,L | 41,L | 40,V | 46,L | 29,L |
| **47** | 42,T | 42,T | 41,D | 47,T | 30,V |
| **48** | 43,N | 43,N | 42,G | 48,G | 31,R |
| **49** | 44,L | 44,L | 43,F | 49,A | 32,P |
| **50** | 45,L | 45,L | 44,L | 50,L | 33,L |

| | | | | | |
|---|---|---|---|---|---|
| **51** | 46,D | 46,D | 45,M | 51,K | 34,R |
| **52** | 47,S | 47,S | 46,G | 52,S | 35,S |
| **53** | 48,D | 48,D | 47,A | 53,D | 36,D |
| **54** | 49,D | 49,D | 48,D | 54,D | 37,D |
| **55** | 50,I | 50,I | 49,N | 55,T | 38,T |
| **56** | 51,R | 51,R | 50,L | 56,R | 39,E |
| **57** | 52,H | 52,H | 51,A | 57,Y | 40,G |
| **58** | 53,M | 53,M | 52,M | 58,M | 41,F |
| **59** | 54,L | 54,L | 53,V | 59,A | 42,A |
| **60** | 55,N | 55,N | 54,S | 60,E | 43,E |
| **61** | 56,A | 56,A | 55,A | 61,A | 44,G |
| **62** | 57,L | 57,L | 56,L | 62,L | 45,L |
| **63** | 58,T | 58,T | 57,Q | 63,R | 46,A |
| **64** | 59,K | 59,K | 58,Q | 64,A | 47,R |
| **65** | 60,L | 60,L | 59,M | 65,M | 48,L |
| **66** | 61,G | 61,G | 60,G | 66,G | 49,G |
| **67** | 62,V | 62,V | 61,A | 67,V | 50,Y |
| **68** | 63,N | 63,N | 62,S | 68,T | 51,R |
| **69** | 64,Y | 64,Y | 63,I | 69,I | 52,V |
| **70** | 65,R | 65,R | 64,Q | 70,D | 53,G |
| **71** | 66,L | 66,L | 65,V | -,- | 54,R |
| **72** | 67,S | 67,S | 66,I | 71,E | 55,T |
| **73** | 68,A | 68,A | 67,E | 72,P | 56,P |
| **74** | 69,D | 69,D | 68,D | 73,D | 57,D |
| **75** | 70,K | 70,K | 69,E | 74,D | -,- |
| **76** | 71,T | 71,T | 70,N | 75,T | -,- |
| **77** | 72,T | 72,T | 71,I | 76,T | 58,S |

| | | | | | |
|---|---|---|---|---|---|
| **78** | 73,C | 73,C | 72,L | 77,F | 59,W |
| **79** | 74,E | 74,E | 73,V | 78,I | 60,Q |
| **80** | 75,V | 75,V | 74,V | 79,V | 61,V |
| **81** | 76,E | 76,E | 75,E | 80,K | 62,D |
| **82** | 77,G | 77,G | 76,G | 81,G | 63,G |
| **83** | 78,L | 78,L | 77,V | 82,S | 64,R |
| **84** | 79,G | 79,G | 78,G | 83,G | 65,P |
| **85** | 80,Q | 80,Q | 79,M | 84,K | 66,Q |
| **86** | 81,A | 81,A | 80,T | 85,L | -,- |
| **87** | 82,F | 82,F | 81,G | 86,Q | 67,G |
| **88** | 83,H | 83,H | 82,L | 87,P | 68,P |
| **89** | 84,T | 84,T | 83,Q | 88,P | 69,A |
| **90** | 85,T | 85,T | 84,A | 89,A | 70,V |
| **91** | 86,Q | 86,Q | 85,P | 90,A | 71,A |
| **92** | 87,P | 87,P | 86,P | 91,P | 72,E |
| **93** | 88,L | 88,L | 87,E | -,- | 73,A |
| **94** | 89,E | 89,E | 88,A | -,- | 74,D |
| **95** | 90,L | 90,L | 89,L | 92,L | 75,V |
| **96** | 91,F | 91,F | 90,D | 93,F | 76,Y |
| **97** | 92,L | 92,L | 91,C | 94,L | 77,C |
| **98** | 93,G | 93,G | 92,G | 95,G | 78,R |
| **99** | 94,N | 94,N | 93,N | 96,N | 79,D |
| **100** | 95,A | 95,A | 94,S | 97,A | 80,G |
| **101** | 96,G | 96,G | 95,G | 98,G | 81,A |
| **102** | 97,T | 97,T | 96,T | 99,T | 82,T |
| **103** | 98,A | 98,A | 97,A | 100,A | 83,T |
| **104** | 99,M | 99,M | 98,I | 101,T | 84,A |

| | | | | | |
|---|---|---|---|---|---|
| **105** | 100,R | 100,R | 99,R | 102,R | 85,R |
| **106** | 101,P | 101,P | 100,L | 103,F | 86,F |
| **107** | 102,L | 102,L | 101,L | 104,L | 87,L |
| **108** | 103,A | 103,A | 102,S | 105,T | 88,P |
| **109** | 104,A | 104,A | 103,G | 106,A | 89,T |
| **110** | 105,A | 105,A | 104,L | 107,A | 90,L |
| **111** | 106,L | 106,L | 105,L | 108,A | 91,A |
| **112** | 107,C | 107,C | 106,A | 109,A | 92,A |
| **113** | 108,L | 108,L | 107,G | 110,L | 93,A |
| **114** | 109,G | 109,G | 108,Q | 111,V | 94,G |
| **115** | 110,Q | 110,Q | 109,P | 112,D | 95,H |
| **116** | 111,G | 111,G | 110,F | 113,G | 96,G |
| **117** | 112,D | 112,D | 111,N | 114,K | 97,T |
| **118** | 113,Y | 113,Y | 112,T | 115,V | 98,Y |
| **119** | 114,V | 114,V | 113,V | 116,I | 99,R |
| **120** | 115,L | 115,L | 114,L | 117,V | 100,F |
| **121** | 116,T | 116,T | 115,T | 118,D | 101,D |
| **122** | 117,G | 117,G | 116,G | 119,G | 102,A |
| **123** | 118,E | 118,E | 117,D | 120,D | 103,S |
| **124** | 119,P | 119,P | 118,S | 121,A | 104,E |
| **125** | 120,R | 120,R | 119,S | 122,H | 105,Q |
| **126** | 121,M | 121,M | 120,L | 123,M | 106,M |
| **127** | 122,K | 122,K | 121,Q | 124,R | 107,R |
| **128** | 123,E | 123,E | 122,R | 125,K | 108,R |
| **129** | 124,R | 124,R | 123,R | 126,R | 109,R |
| **130** | 125,P | 125,P | 124,P | 127,P | 110,P |
| **131** | 126,I | 126,I | 125,M | 128,I | 111,L |

| | | | | | |
|---|---|---|---|---|---|
| **132** | 127,G | 127,G | 126,K | 129,G | 112,L |
| **133** | 128,H | 128,H | 127,R | 130,P | 113,P |
| **134** | 129,L | 129,L | 128,I | 131,L | 114,L |
| **135** | 130,V | 130,V | 129,I | 132,V | 115,T |
| **136** | 131,D | 131,D | 130,D | 133,D | 116,R |
| **137** | 132,A | 132,A | 131,P | 134,A | 117,A |
| **138** | 133,L | 133,L | 132,L | 135,L | 118,L |
| **139** | 134,R | 134,R | 133,T | 136,R | 119,R |
| **140** | 135,Q | 135,Q | 134,L | 137,S | 120,E |
| **141** | 136,A | 136,A | 135,M | 138,L | 121,L |
| **142** | 137,G | 137,G | 136,G | 139,G | 122,G |
| **143** | 138,A | 138,A | 137,A | 140,I | 123,V |
| **144** | 139,Q | 139,Q | 138,K | 141,D | 124,D |
| **145** | 140,I | 140,I | 139,I | 142,A | 125,L |
| **146** | 141,E | 141,E | 140,D | 143,S | 126,R |
| **147** | 142,Y | 142,Y | 141,S | 144,A | 127,H |
| **148** | 143,L | 143,L | 142,T | 145,E | 128,E |
| **149** | 144,E | 144,E | 143,G | 146,T | 129,E |
| **150** | 145,Q | 145,Q | -,- | -,- | 130,R |
| **151** | 146,E | 146,E | -,- | -,- | 131,D |
| **152** | 147,N | 147,N | 144,N | 147,G | 132,G |
| **153** | 148,F | 148,F | 145,V | 148,C | 133,H |
| **154** | 149,P | 149,P | 146,P | 149,P | 134,H |
| **155** | 150,P | 150,P | 147,P | 150,P | 135,P |
| **156** | 151,L | 151,L | 148,L | 151,V | 136,L |
| **157** | 152,R | 152,R | 149,K | 152,T | 137,T |
| **158** | 153,I | 153,I | 150,I | 153,I | 138,V |

| | | | | | |
|---|---|---|---|---|---|
| **159** | 154,Q | 154,Q | 151,Y | 154,N | 139,R |
| **160** | 155,G | 155,G | 152,G | 155,G | 140,A |
| **161** | 156,T | 156,T | 153,N | 156,T | 141,A |
| **162** | 157,G | 157,G | 154,P | 157,G | 142,G |
| **163** | -,- | -,- | 155,R | 158,R | -,- |
| **164** | 158,L | 158,L | 156,L | 159,F | 143,V |
| **165** | 159,Q | 159,Q | 157,T | 160,E | 144,A |
| **166** | 160,A | 160,A | 158,G | 161,A | 145,G |
| **167** | 161,G | 161,G | 159,I | 162,S | 146,G |
| **168** | 162,T | 162,T | 160,H | 163,R | 147,E |
| **169** | 163,V | 163,V | 161,Y | 164,V | 148,V |
| **170** | 164,T | 164,T | 162,Q | 165,Q | 149,T |
| **171** | 165,I | 165,I | 163,L | 166,I | 150,L |
| **172** | 166,D | 166,D | 164,P | 167,D | 151,D |
| **173** | 167,G | 167,G | 165,M | 168,G | 152,A |
| **174** | 168,S | 168,S | 166,A | 169,G | 153,G |
| **175** | 169,I | 169,I | -,- | 170,L | 154,Q |
| **176** | 170,S | 170,S | 167,S | 171,S | 155,S |
| **177** | 171,S | 171,S | 168,A | 172,S | 156,S |
| **178** | 172,Q | 172,Q | 169,Q | 173,Q | 157,Q |
| **179** | 173,F | 173,F | 170,V | 174,Y | 158,Y |
| **180** | 174,L | 174,L | 171,K | 175,V | 159,L |
| **181** | 175,T | 175,T | 172,S | 176,S | 160,T |
| **182** | 176,A | 176,A | 173,C | 177,A | 161,A |
| **183** | 177,F | 177,F | 174,L | 178,L | 162,L |
| **184** | 178,L | 178,L | 175,L | 179,L | 163,L |
| **185** | 179,M | 179,M | 176,L | 180,M | 164,L |
| **186** | 180,S | 180,S | 177,A | 181,M | 165,L |

| | | | | | |
|---|---|---|---|---|---|
| **187** | 181,A | 181,A | 178,G | 182,A | 166,G |
| **188** | 182,P | 182,P | 179,L | 183,A | 167,P |
| **189** | 183,L | 183,L | 180,Y | 184,G | 168,L |
| **190** | 184,A | 184,A | 181,A | 185,G | 169,T |
| **191** | 185,Q | 185,Q | 182,R | 186,D | 170,E |
| **192** | 186,G | 186,G | 183,G | 187,R | 171,K |
| **193** | 187,K | 187,K | 184,K | 188,A | 172,G |
| **194** | 188,V | 188,V | 185,T | 189,V | 173,L |
| **195** | 189,T | 189,T | 186,C | 190,D | 174,R |
| **196** | 190,I | 190,I | 187,I | 191,V | 175,I |
| **197** | 191,K | 191,K | 188,T | 192,E | 176,H |
| **198** | 192,I | 192,I | 189,E | 193,L | 177,V |
| **199** | 193,V | 193,V | -,- | 194,L | 178,T |
| **200** | 194,G | 194,G | -,- | 195,G | -,- |
| **201** | 195,E | 195,E | -,- | 196,E | 179,D |
| **202** | 196,L | 196,L | -,- | 197,H | 180,L |
| **203** | 197,V | 197,V | -,- | 198,I | 181,V |
| **204** | -,- | -,- | -,- | 199,G | -,- |
| **205** | 198,S | 198,S | 190,P | 200,A | 182,S |
| **206** | 199,K | 199,K | 191,A | 201,L | 183,V |
| **207** | 200,P | 200,P | 192,P | 202,G | 184,P |
| **208** | 201,Y | 201,Y | 193,S | 203,Y | 185,Y |
| **209** | 202,I | 202,I | 194,R | 204,I | 186,I |
| **210** | 203,D | 203,D | 195,D | 205,D | 187,E |
| **211** | 204,I | 204,I | 196,H | 206,L | 188,I |
| **212** | 205,T | 205,T | 197,T | 207,T | 189,T |
| **213** | 206,L | 206,L | 198,E | 208,V | 190,L |

| | | | | | |
|---|---|---|---|---|---|
| **214** | 207,H | 207,H | 199,R | 209,A | 191,A |
| **215** | 208,I | 208,I | 200,L | 210,A | 192,M |
| **216** | 209,M | 209,M | 201,L | 211,M | 193,M |
| **217** | 210,E | 210,E | 202,K | 212,R | 194,R |
| **218** | 211,Q | 211,Q | 203,H | 213,A | 195,A |
| **219** | 212,F | 212,F | 204,F | 214,F | 196,F |
| **220** | 213,G | 213,G | 205,H | 215,G | 197,G |
| **221** | 214,V | 214,V | 206,Y | 216,A | 198,V |
| **222** | 215,Q | 215,Q | 207,T | 217,K | 199,E |
| **223** | 216,V | 216,V | 208,L | 218,V | 200,V |
| **224** | 217,I | 217,I | 209,Q | 219,E | 201,T |
| **225** | 218,N | 218,N | 210,K | 220,R | 202,R |
| **226** | 219,H | 219,H | 211,D | 221,V | 203,E |
| **227** | 220,D | 220,D | 212,K | 222,S | 204,G |
| **228** | 221,Y | 221,Y | 213,Q | 223,P | 205,H |
| **229** | 222,Q | 222,Q | -,- | 224,V | -,- |
| **230** | 223,E | 223,E | 214,S | 225,A | 206,D |
| **231** | 224,F | 224,F | 215,I | 226,W | 207,F |
| **232** | 225,V | 225,V | 216,C | 227,R | 208,V |
| **233** | 226,I | 226,I | 217,V | 228,V | 209,V |
| **234** | 227,P | 227,P | 218,S | 229,E | 210,P |
| **235** | 228,A | 228,A | 219,G | 230,P | 211,P |
| **236** | 229,G | 229,G | 220,G | 231,T | 212,G |
| **237** | 230,Q | 230,Q | 221,G | 232,G | 213,G |
| **238** | 231,S | 231,S | 222,K | -,- | -,- |
| **239** | 232,Y | 232,Y | 223,L | 233,Y | 214,Y |
| **240** | 233,V | 233,V | -,- | -,- | -,- |

| | | | | | |
|---|---|---|---|---|---|
| **241** | 234,S | 234,S | 224,K | 234,H | 215,R |
| **242** | 235,P | 235,P | 225,A | 235,A | 216,A |
| **243** | 236,G | 236,G | 226,N | 236,A | 217,T |
| **244** | 237,Q | 237,Q | 227,D | 237,D | 218,T |
| **245** | 238,F | 238,F | 228,I | 238,F | 219,Y |
| **246** | 239,L | 239,L | 229,S | 239,V | 220,A |
| **247** | 240,V | 240,V | 230,I | 240,I | 221,I |
| **248** | 241,E | 241,E | 231,P | 241,E | 222,E |
| **249** | 242,G | 242,G | 232,G | 242,P | 223,P |
| **250** | 243,D | 243,D | 233,D | 243,D | 224,D |
| **251** | 244,A | 244,A | 234,I | 244,A | 225,A |
| **252** | 245,S | 245,S | 235,S | 245,S | 226,S |
| **253** | 246,S | 246,S | 236,S | 246,A | 227,T |
| **254** | 247,A | 247,A | 237,A | 247,A | 228,S |
| **255** | 248,S | 248,S | 238,A | 248,T | 229,S |
| **256** | 249,Y | 249,Y | 239,F | 249,Y | 230,Y |
| **257** | 250,F | 250,F | 240,F | 250,L | 231,F |
| **258** | 251,L | 251,L | 241,I | 251,W | 232,F |
| **259** | 252,A | 252,A | 242,V | 252,A | 233,A |
| **260** | 253,A | 253,A | 243,A | 253,A | 234,A |
| **261** | 254,A | 254,A | 244,A | 254,E | 235,A |
| **262** | 255,A | 255,A | 245,T | 255,V | 236,A |
| **263** | 256,I | 256,I | 246,I | 256,L | 237,L |
| **264** | 257,K | 257,K | 247,T | 257,S | 238,S |
| **265** | -,- | -,- | 248,P | -,- | -,- |
| **266** | 258,G | 258,G | 249,G | 258,G | 239,G |
| **267** | 259,G | 259,G | 250,S | 259,G | 240,G |

| | | | | | |
|---|---|---|---|---|---|
| **268** | 260,E | 260,E | 251,A | 260,K | 241,E |
| **269** | 261,V | 261,V | 252,I | 261,I | 242,V |
| **270** | 262,K | 262,K | 253,R | 262,D | 243,T |
| **271** | 263,V | 263,V | 254,L | 263,L | 244,V |
| **272** | 264,T | 264,T | 255,C | 264,G | 245,P |
| **273** | 265,G | 265,G | 256,R | 265,T | 246,G |
| **274** | 266,I | 266,I | 257,V | 266,P | 247,L |
| **275** | 267,G | 267,G | 258,G | 267,A | 248,G |
| **276** | 268,K | 268,K | 259,V | 268,E | 249,E |
| **277** | 269,N | 269,N | 260,N | 269,Q | 250,G |
| **278** | 270,S | 270,S | 261,P | 270,F | 251,A |
| **279** | 271,I | 271,I | 262,T | 271,S | 252,L |
| **280** | 272,Q | 272,Q | 263,R | 272,Q | 253,Q |
| **281** | 273,G | 273,G | -,- | 273,P | 254,G |
| **282** | 274,D | 274,D | -,- | 274,D | 255,D |
| **283** | 275,I | 275,I | 264,L | 275,A | 256,L |
| **284** | 276,Q | 276,Q | 265,G | 276,K | 257,G |
| **285** | 277,F | 277,F | 266,V | 277,A | 258,F |
| **286** | 278,A | 278,A | 267,I | 278,Y | 259,V |
| **287** | 279,D | 279,D | 268,N | 279,D | 260,D |
| **288** | 280,A | 280,A | 269,L | 280,L | 261,V |
| **289** | 281,L | 281,L | 270,L | 281,I | 262,L |
| **290** | 282,E | 282,E | 271,K | 282,S | 263,R |
| **291** | 283,K | 283,K | 272,M | 283,K | 264,R |
| **292** | 284,M | 284,M | 273,M | -,- | 265,M |
| **293** | 285,G | 285,G | 274,G | -,- | 266,G |
| **294** | 286,A | 286,A | 275,A | -,- | 267,A |

| | | | | | |
|---|---|---|---|---|---|
| **295** | 287,Q | 287,Q | 276,D | -,- | 268,E |
| **296** | 288,I | 288,I | 277,I | -,- | 269,V |
| **297** | 289,E | 289,E | 278,E | -,- | 270,E |
| **298** | 290,W | 290,W | 279,V | -,- | 271,I |
| **299** | 291,G | 291,G | 280,T | -,- | 272,G |
| **300** | -,- | -,- | 281,H | -,- | -,- |
| **301** | -,- | -,- | 282,Y | -,- | -,- |
| **302** | -,- | -,- | 283,T | -,- | -,- |
| **303** | -,- | -,- | 284,E | -,- | -,- |
| **304** | -,- | -,- | 285,K | -,- | -,- |
| **305** | -,- | -,- | 286,N | -,- | -,- |
| **306** | -,- | -,- | 287,E | -,- | -,- |
| **307** | -,- | -,- | 288,E | -,- | -,- |
| **308** | -,- | -,- | 289,P | -,- | -,- |
| **309** | 292,D | 292,D | 290,T | -,- | 273,A |
| **310** | 293,D | 293,D | 291,A | -,- | 274,D |
| **311** | 294,Y | 294,Y | 292,D | -,- | 275,R |
| **312** | 295,V | 295,V | 293,I | -,- | 276,T |
| **313** | 296,I | 296,I | 294,T | -,- | 277,T |
| **314** | 297,A | 297,A | 295,V | -,- | 278,V |
| **315** | 298,R | 298,R | 296,R | -,- | 279,R |
| **316** | -,- | -,- | -,- | -,- | 280,G |
| **317** | 299,R | 299,R | 297,H | 284,F | 281,T |
| **318** | 300,G | 300,G | 298,A | 285,P | 282,G |
| **319** | 301,E | 301,E | 299,R | 286,H | 283,E |
| **320** | 302,L | 302,L | 300,L | 287,L | 284,L |
| **321** | 303,N | 303,N | 301,K | 288,P | 285,R |
| **322** | 304,A | 304,A | 302,G | 289,A | 286,G |

| | | | | | |
|---|---|---|---|---|---|
| **323** | 305,V | 305,V | 303,I | 290,V | 287,L |
| **324** | 306,D | 306,D | 304,D | -,- | 288,T |
| **325** | 307,L | 307,L | 305,I | 291,I | 289,V |
| **326** | 308,D | 308,D | 306,P | 292,D | 290,N |
| **327** | 309,F | 309,F | 307,P | 293,G | 291,M |
| **328** | 310,N | 310,N | 308,D | 294,S | 292,R |
| **329** | 311,H | 311,H | 309,Q | 295,Q | 293,D |
| **330** | 312,I | 312,I | 310,V | 296,M | 294,I |
| **331** | 313,P | 313,P | 311,P | 297,Q | 295,S |
| **332** | -,- | -,- | 312,L | -,- | -,- |
| **333** | -,- | -,- | 313,T | -,- | -,- |
| **334** | -,- | -,- | 314,I | -,- | -,- |
| **335** | 314,D | 314,D | 315,D | 298,D | 296,D |
| **336** | 315,A | 315,A | 316,E | 299,A | 297,T |
| **337** | 316,A | 316,A | 317,F | 300,I | 298,M |
| **338** | 317,M | 317,M | 318,P | 301,P | 299,P |
| **339** | 318,T | 318,T | 319,V | 302,T | 300,T |
| **340** | 319,I | 319,I | 320,L | 303,L | 301,L |
| **341** | 320,A | 320,A | 321,L | 304,A | 302,A |
| **342** | 321,T | 321,T | 322,I | 305,V | 303,A |
| **343** | 322,T | 322,T | 323,A | 306,L | 304,I |
| **344** | 323,A | 323,A | 324,A | 307,A | 305,A |
| **345** | 324,L | 324,L | 325,A | 308,A | 306,P |
| **346** | 325,F | 325,F | 326,V | 309,F | 307,F |
| **347** | 326,A | 326,A | 327,A | 310,N | 308,A |
| **348** | 327,K | 327,K | 328,Q | 311,E | 309,S |
| **349** | 328,G | 328,G | 329,G | 312,M | 310,G |

| | | | | | |
|---|---|---|---|---|---|
| **350** | 329,T | 329,T | 330,K | 313,P | 311,P |
| **351** | 330,T | 330,T | 331,T | 314,V | 312,V |
| **352** | 331,A | 331,A | 332,V | 315,R | 313,R |
| **353** | 332,I | 332,I | 333,L | 316,F | 314,I |
| **354** | 333,R | 333,R | 334,R | 317,V | 315,E |
| **355** | 334,N | 334,N | 335,D | 318,G | 316,D |
| **356** | 335,V | 335,V | 336,A | 319,I | 317,V |
| **357** | 336,Y | 336,Y | 337,A | 320,E | 318,A |
| **358** | 337,N | 337,N | 338,E | 321,N | 319,N |
| **359** | 338,W | 338,W | 339,L | 322,L | 320,T |
| **360** | 339,R | 339,R | 340,R | 323,R | 321,R |
| **361** | 340,V | 340,V | 341,V | 324,V | 322,V |
| **362** | 341,K | 341,K | 342,K | 325,K | 323,K |
| **363** | 342,E | 342,E | 343,E | 326,E | 324,E |
| **364** | 343,T | 343,T | 344,T | 327,C | 325,C |
| **365** | 344,D | 344,D | 345,D | 328,D | 326,D |
| **366** | 345,R | 345,R | 346,R | 329,R | 327,R |
| **367** | 346,L | 346,L | 347,I | 330,I | 328,L |
| **368** | 347,A | 347,A | 348,A | 331,R | 329,E |
| **369** | 348,A | 348,A | 349,A | 332,A | 330,A |
| **370** | 349,M | 349,M | 350,M | 333,L | 331,C |
| **371** | 350,A | 350,A | 351,V | 334,S | 332,A |
| **372** | 351,T | 351,T | 352,D | 335,S | 333,E |
| **373** | 352,E | 352,E | 353,G | 336,G | 334,N |
| **374** | 353,L | 353,L | 354,L | 337,L | 335,L |
| **375** | 354,R | 354,R | 355,Q | 338,S | 336,R |
| **376** | 355,K | 355,K | 356,K | 339,R | 337,R |

| | | | | | |
|---|---|---|---|---|---|
| **377** | 356,V | 356,V | 357,L | 340,I | 338,L |
| **378** | -,- | -,- | -,- | 341,V | -,- |
| **379** | -,- | -,- | -,- | 342,P | -,- |
| **380** | 357,G | 357,G | 358,G | 343,N | 339,G |
| **381** | 358,A | 358,A | 359,I | 344,L | 340,V |
| **382** | 359,T | 359,T | 360,A | 345,G | 341,R |
| **383** | 360,V | 360,V | 361,A | 346,T | 342,V |
| **384** | 361,E | 361,E | 362,E | 347,E | 343,E |
| **385** | 362,E | 362,E | 363,S | 348,E | 344,T |
| **386** | 363,G | 363,G | 364,L | 349,G | 345,G |
| **387** | 364,E | 364,E | 365,P | 350,D | 346,P |
| **388** | 365,D | 365,D | 366,D | 351,D | 347,D |
| **389** | 366,F | 366,F | 367,G | 352,L | 348,W |
| **390** | 367,I | 367,I | 368,V | 353,I | 349,I |
| **391** | 368,V | 368,V | 369,I | 354,I | 350,E |
| **392** | 369,I | 369,I | 370,I | 355,A | 351,I |
| **393** | 370,T | 370,T | 371,Q | 356,S | 352,H |
| **394** | 371,P | 371,P | 372,G | 357,D | 353,P |
| **395** | 372,P | 372,P | 373,G | 358,P | 354,G |
| **396** | 373,T | 373,T | 374,T | 359,S | 355,A |
| **397** | -,- | -,- | -,- | 360,L | -,- |
| **398** | -,- | -,- | -,- | 361,A | -,- |
| **399** | -,- | -,- | -,- | 362,G | -,- |
| **400** | 374,K | 374,K | -,- | 363,K | 356,T |
| **401** | 375,L | 375,L | 375,L | 364,I | 357,P |
| **402** | 376,I | 376,I | 376,E | 365,L | 358,T |
| **403** | 377,H | 377,H | 377,G | 366,T | 359,G |

| | | | | | |
|---|---|---|---|---|---|
| **404** | 378,A | 378,A | 378,G | 367,A | 360,A |
| **405** | 379,A | 379,A | 379,E | 368,E | 361,E |
| **406** | 380,I | 380,I | 380,V | 369,I | 362,I |
| **407** | 381,D | 381,D | 381,N | 370,D | 363,K |
| **408** | 382,T | 382,T | 382,S | 371,S | 364,T |
| **409** | 383,Y | 383,Y | 383,Y | 372,F | 365,Y |
| **410** | 384,D | 384,D | 384,D | 373,A | 366,G |
| **411** | 385,D | 385,D | 385,D | 374,D | 367,D |
| **412** | 386,H | 386,H | 386,H | 375,H | 368,H |
| **413** | 387,R | 387,R | 387,R | 376,R | 369,R |
| **414** | 388,M | 388,M | 388,I | 377,I | 370,I |
| **415** | 389,A | 389,A | 389,A | 378,A | 371,V |
| **416** | 390,M | 390,M | 390,M | 379,M | 372,M |
| **417** | 391,C | 391,C | 391,A | 380,S | 373,S |
| **418** | 392,F | 392,F | 392,F | 381,F | 374,F |
| **419** | 393,S | 393,S | 393,A | 382,A | 375,A |
| **420** | 394,L | 394,L | 394,V | 383,L | 376,V |
| **421** | 395,V | 395,V | 395,A | 384,A | 377,T |
| **422** | 396,A | 396,A | 396,G | 385,G | 378,G |
| **423** | -,- | -,- | 397,T | -,- | -,- |
| **424** | 397,L | 397,L | 398,L | 386,L | 379,L |
| **425** | 398,S | 398,S | 399,A | 387,K | 380,R |
| **426** | 399,D | 399,D | 400,K | 388,I | 381,V |
| **427** | 400,T | 400,T | 401,G | 389,G | 382,P |
| **428** | 401,P | 401,P | 402,P | 390,G | 383,G |
| **429** | 402,V | 402,V | 403,V | 391,I | 384,I |
| **430** | 403,T | 403,T | 404,R | 392,T | 385,S |

| | | | | | |
|---|---|---|---|---|---|
| **431** | 404,I | 404,I | 405,I | 393,I | 386,F |
| **432** | 405,N | 405,N | 406,R | 394,L | 387,D |
| **433** | 406,D | 406,D | 407,N | 395,D | 388,D |
| **434** | 407,P | 407,P | 408,C | 396,P | 389,P |
| **435** | 408,K | 408,K | 409,D | 397,D | 390,G |
| **436** | 409,C | 409,C | 410,N | 398,C | 391,C |
| **437** | 410,T | 410,T | 411,V | 399,V | 392,V |
| **438** | 411,S | 411,S | 412,K | 400,A | 393,R |
| **439** | 412,K | 412,K | 413,T | 401,K | 394,K |
| **440** | 413,T | 413,T | 414,S | 402,T | 395,T |
| **441** | 414,F | 414,F | 415,F | 403,F | 396,F |
| **442** | 415,P | 415,P | 416,P | 404,P | 397,P |
| **443** | 416,D | 416,D | 417,N | 405,S | 398,G |
| **444** | 417,Y | 417,Y | 418,F | 406,Y | 399,F |
| **445** | 418,F | 418,F | 419,V | 407,W | 400,H |
| **446** | 419,D | 419,D | 420,E | 408,N | 401,E |
| **447** | 420,K | 420,K | 421,L | 409,V | 402,E |
| **448** | 421,F | 421,F | 422,A | 410,L | 403,F |
| **449** | 422,A | 422,A | 423,N | 411,S | 404,G |
| **450** | 423,Q | 423,Q | 424,E | 412,S | 405,A |
| **451** | 424,L | 424,L | 425,V | 413,L | 406,L |
| **452** | 425,S | 425,S | 426,G | 414,G | 407,R |
| **453** | 426,R | 426,R | 427,M | 415,V | 408,A |
| **454** | -,- | -,- | 428,N | 416,A | 409,R |
| **455** | -,- | -,- | 429,V | 417,Y | 410,L |
| **456** | -,- | -,- | 430,K | 418,E | -,- |
| **457** | -,- | -,- | 431,G | 419,D | -,- |

| | | | | | |
|---|---|---|---|---|---|
| **458** | -,- | -,- | 432,V | -,- | -,- |
| **459** | -,- | -,- | 433,R | -,- | -,- |
| **460** | -,- | -,- | 434,G | -,- | -,- |
| **461** | -,- | -,- | 435,R | -,- | -,- |
| **462** | -,- | -,- | 436,G | -,- | -,- |
| **463** | -,- | -,- | 437,G | -,- | -,- |
| **464** | -,- | -,- | 438,F | -,- | -,- |

**Supplementary table 2. Class I-IV EPSPS reference sequences**

```
>Vibrio cholerae|vbEPSPS
MESLTLQPIELISGEVNLPGSKSVSNRALLLAALASGTTRLTNLLDSDDIRHMLNALTKL
GVNYRLSADKTTCEVEGLGQAFHTTQPLELFLGNAGTAMRPLAAALCLGQGDYVLTGEPR
MKERPIGHLVDALRQAGAQIEYLEQENFPPLRIQGTGLQAGTVTIDGSISSQFLTAFLMS
APLAQGKVTIKIVGELVSKPYIDITLHIMEQFGVQVINHDYQEFVIPAGQSYVSPGQFLV
EGDASSASYFLAAAAIKGGEVKVTGIGKNSIQGDIQFADALEKMGAQIEWGDDYVIARRG
ELNAVDLDFNHIPDAAMTIATTALFAKGTTAIRNVYNWRVKETDRLAAMATELRKVGATV
EEGEDFIVITPPTKLIHAAIDTYDDHRMAMCFSLVALSDTPVTINDPKCTSKTFPDYFDK
FAQLSR
```

```
>Coxiella burnetii|cbESPSP
MDYQTIPSQGLSGEICVPGDKSISHRAVLLAAIAEGQTQVDGFLMGADNLAMVSALQQMG
ASIQVIEDENILVVEGVGMTGLQAPPEALDCGNSGTAIRLLSGLLAGQPFNTVLTGDSSL
QRRPMKRIIDPLTLMGAKIDSTGNVPPLKIYGNPRLTGIHYQLPMASAQVKSCLLLAGLY
ARGKTCITEPAPSRDHTERLLKHFHYTLQKDKQSICVSGGGKLKANDISIPGDISSAAFF
IVAATITPGSAIRLCRVGVNPTRLGVINLLKMMGADIEVTHYTEKNEEPTADITVRHARL
KGIDIPPDQVPLTIDEFPVLLIAAAVAQGKTVLRDAAELRVKETDRIAAMVDGLQKLGIA
AESLPDGVIIQGGTLEGGEVNSYDDHRIAMAFAVAGTLAKGPVRIRNCDNVKTSFPNFVE
LANEVGMNVKGVRGRGGF
```

```
>Brevundimonas vesicularis|bvEPSPS
MMMGRAKLTIIPPGKPLTGRAMPPGSKSITNRALLLAGLAKGTSRLTGALKSDDTRYMAE
ALRAMGVTIDEPDDTTFIVKGSGKLQPPAAPLFLGNAGTATRFLTAAAALVDGKVIVDGD
AHMRKRPIGPLVDALRSLGIDASAETGCPPVTINGTGRFEASRVQIDGGLSSQYVSALLM
MAAGGDRAVDVELLGEHIGALGYIDLTVAAMRAFGAKVERVSPVAWRVEPTGYHAADFVI
EPDASAATYLWAAEVLSGGKIDLGTPAEQFSQPDAKAYDLISKFPHLPAVIDGSQMQDAI
PTLAVLAAFNEMPVRFVGIENLRVKECDRIRALSSGLSRIVPNLGTEEGDDLIIASDPSL
AGKILTAEIDSFADHRIAMSFALAGLKIGGITILDPDCVAKTFPSYWNVLSSLGVAYED
```

```
>Streptomyces davawensis|sdEPSPS
MPVADIPGSKSITARALFLAAAADGVTTLVRPLRSDDTEGFAEGLARLGYRVGRTPDSWQ
VDGRPQGPAVAEADVYCRDGATTARFLPTLAAAGHGTYRFDASEQMRRRPLLPLTRALRE
LGVDLRHEERDGHHPLTVRAAGVAGGEVTLDAGQSSQYLTALLLLGPLTEKGLRIHVTDL
VSVPYIEITLAMMRAFGVEVTREGHDFVVPPGGYRATTYAIEPDASTSSYFFAAAALSGG
EVTVPGLGEGALQGDLGFVDVLRRMGAEVEIGADRTTVRGTGELRGLTVNMRDISDTMPT
LAAIAPFASGPVRIEDVANTRVKECDRLEACAENLRRLGVRVETGPDWIEIHPGATPTGA
EIKTYGDHRIVMSFAVTGLRVPGISFDDPGCVRKTFPGFHEEFGALRARL
```

**Supplementary table 3 . Taxonomic distribution of EPSPS proteins by class.** EPSPS sequences were obtained from Pfam database [1] .

| Taxonomy | Class Iα | Class Iβ | Class II | Class III | Class IV | None |
|---|---|---|---|---|---|---|
| **Archaea** | 15 | 142 | 0 | 4 | 0 | 35 |
| **Bacteria** | 1511 | 1460 | 1376 | 217 | 39 | 5447 |
| **Eukaryota** | 469 | 488 | 0 | 7 | 0 | 123 |
| **Acidobacteria** | 1 | 1 | 10 | 0 | 0 | 13 |
| **Actinobacteria** | 490 | 79 | 17 | 16 | 34 | 953 |
| **Alveolata** | 6 | 10 | 0 | 0 | 0 | 25 |
| **Amoebozoa** | 1 | 1 | 0 | 0 | 0 | 0 |
| **Aquificae** | 0 | 0 | 12 | 0 | 0 | 14 |
| **Armatimonadetes** | 2 | 0 | 1 | 1 | 0 | 5 |
| **Asgard group** | 0 | 1 | 0 | 0 | 0 | 0 |
| **Bacteroidetes** | 372 | 419 | 4 | 20 | 3 | 421 |
| **Caldiserica** | 0 | 0 | 0 | 0 | 0 | 1 |
| **Calditrichaeota** | 0 | 0 | 0 | 0 | 0 | 3 |
| **Chlamydiae** | 1 | 9 | 0 | 0 | 0 | 19 |
| **Chlorobi** | 0 | 0 | 11 | 0 | 0 | 15 |
| **Chloroflexi** | 4 | 10 | 6 | 3 | 0 | 45 |
| **Choanoflagellida** | 2 | 2 | 0 | 0 | 0 | 0 |
| **Chrysiogenetes** | 0 | 0 | 1 | 0 | 0 | 1 |
| **Crenarchaeota** | 0 | 21 | 0 | 2 | 0 | 9 |
| **Cryptophyta** | 1 | 1 | 0 | 0 | 0 | 0 |
| **Cyanobacteria** | 8 | 8 | 98 | 1 | 0 | 118 |
| **Deferribacteres** | 0 | 0 | 5 | 1 | 0 | 5 |
| **Deinococcus Thermus** | 2 | 12 | 3 | 1 | 2 | 17 |
| **Dictyoglomi** | 0 | 0 | 1 | 0 | 0 | 1 |
| **Elusimicrobia** | 0 | 0 | 1 | 0 | 0 | 4 |

| | | | | | | |
|---|---|---|---|---|---|---|
| **Environmental samples** | 0 | 4 | 0 | 0 | 0 | 0 |
| **Euryarchaeota** | 14 | 99 | 0 | 1 | 0 | 27 |
| **Fibrobacteres** | 1 | 3 | 0 | 0 | 0 | 5 |
| **Firmicutes** | 28 | 236 | 440 | 21 | 0 | 1909 |
| **Fungi** | 358 | 368 | 0 | 6 | 0 | 57 |
| **Fusobacteria** | 1 | 14 | 0 | 0 | 0 | 21 |
| **Gemmatimonadetes** | 1 | 1 | 3 | 0 | 0 | 12 |
| **Haloplasmatales** | 0 | 0 | 2 | 0 | 0 | 1 |
| **Haptophyceae** | 0 | 0 | 0 | 0 | 0 | 4 |
| **Ichthyosporea** | 2 | 2 | 0 | 0 | 0 | 0 |
| **Ignavibacteriae** | 0 | 0 | 2 | 0 | 0 | 2 |
| **Kiritimatiellaeota** | 0 | 0 | 1 | 0 | 0 | 1 |
| **Lentisphaerae** | 1 | 1 | 0 | 0 | 0 | 1 |
| **Metazoa** | 0 | 0 | 0 | 0 | 0 | 8 |
| **Nitrospinae Tectomicrobia group** | 1 | 1 | 0 | 1 | 0 | 6 |
| **Nitrospirae** | 1 | 2 | 11 | 0 | 0 | 22 |
| **Nucleariidae & Fonticula group** | 0 | 1 | 0 | 0 | 0 | 0 |
| **Planctomycetes** | 20 | 22 | 0 | 8 | 0 | 17 |
| **Proteobacteria** | 549 | 592 | 714 | 149 | 0 | 1705 |
| **Rhizaria** | 1 | 1 | 0 | 1 | 0 | 0 |
| **Rhodophyta** | 2 | 2 | 0 | 0 | 0 | 0 |
| **Spirochaetes** | 2 | 18 | 4 | 1 | 0 | 61 |
| **Stramenopiles** | 22 | 23 | 0 | 0 | 0 | 3 |
| **Synergistetes** | 0 | 4 | 7 | 0 | 0 | 14 |
| **Tenericutes** | 0 | 4 | 0 | 0 | 0 | 7 |
| **Thaumarchaeota** | 0 | 7 | 0 | 0 | 0 | 5 |
| **Thermobaculum** | 0 | 0 | 1 | 1 | 0 | 1 |
| **Thermodesulfobacteria** | 1 | 2 | 0 | 1 | 0 | 4 |

| | | | | | | |
|---|---|---|---|---|---|---|
| **Thermotogae** | 0 | 2 | 5 | 0 | 0 | 15 |
| **Unclassified Parcubacteria group** | 2 | 3 | 0 | 0 | 0 | 100 |
| **Verrucomicrobia** | 10 | 1 | 7 | 1 | 0 | 19 |
| **Viridiplantae** | 74 | 77 | 0 | 0 | 0 | 68 |

**Supplementary table 4 . Taxonomic distribution of EPSPS proteins in archaea and bacteria species by Class.** EPSPS sequences were obtained from the COG database [2].

| Taxonomy | Class Iα | Class Iβ | Class II | Class III | Class IV | None |
|---|---|---|---|---|---|---|
| **Archaea** | **5** | **63** | **0** | **2** | **0** | **13** |
| **Crenarchaeota** | 0 | 10 | 0 | 1 | 0 | 5 |
| **Euryarchaeota** | 5 | 50 | 0 | 1 | 0 | 6 |
| **Korarchaeota** | 0 | 0 | 0 | 0 | 0 | 1 |
| **Thaumarchaeota** | 0 | 3 | 0 | 0 | 0 | 1 |
| **Bacteria** | **234** | **223** | **252** | **26** | **2** | **64** |
| **Acidobacteria** | 0 | 0 | 6 | 0 | 0 | 0 |
| **Actinobacteria** | 0 | 0 | 0 | 0 | 0 | 0 |
| **Aquificae** | 0 | 0 | 8 | 0 | 0 | 0 |
| **Bacteroidetes** | 37 | 44 | 2 | 0 | 0 | 7 |
| **Chlamydiae** | 0 | 5 | 0 | 0 | 0 | 1 |
| **Chlorobi** | 0 | 0 | 0 | 0 | 0 | 0 |
| **Chloroflexi** | 2 | 4 | 3 | 2 | 0 | 2 |
| **Cyanobacteria** | 3 | 3 | 29 | 0 | 0 | 0 |
| **Deinococcus-thermus** | 2 | 2 | 3 | 1 | 0 | 0 |
| **Firmicutes (Bacilli)** | 3 | 2 | 25 | 1 | 0 | 4 |
| **Firmicutes (Clostridia)** | 0 | 13 | 28 | 1 | 0 | 4 |
| **Firmicutes (Mollicutes)** | 0 | 0 | 0 | 0 | 0 | 1 |
| **Firmicutes (Other)** | 0 | 3 | 2 | 0 | 0 | 0 |
| **Fusobacteria** | 0 | 4 | 0 | 0 | 0 | 0 |
| **Other Bacteria** | 6 | 3 | 15 | 2 | 0 | 4 |
| **Planctomycetes** | 6 | 6 | 0 | 3 | 0 | 0 |

| | | | | | | |
|---|---|---|---|---|---|---|
| **Proteobacteria (Alpha)** | 4 | 4 | 57 | 9 | 0 | 7 |
| **Proteobacteria (Beta)** | 47 | 47 | 0 | 2 | 0 | 2 |
| **Proteobacteria (Delta)** | 14 | 17 | 6 | 3 | 0 | 9 |
| **Proteobacteria (Epsilon)** | 1 | 1 | 9 | 0 | 0 | 1 |
| **Proteobacteria (Gamma)** | 52 | 54 | 41 | 0 | 0 | 8 |
| **Spirochaetes** | 0 | 2 | 1 | 1 | 0 | 3 |
| **Synergistetes** | 0 | 1 | 3 | 0 | 0 | 0 |
| **Thermotogae** | 0 | 0 | 0 | 0 | 0 | 0 |

**Supplementary table 5. Distributions of EPSPS and EPSPS-associated domains across archaea, bacteria and eukaryotes.** Pfam [1] codes of the domains: PF00275 (EPSP_synthase); PF01264 (Chorismate_synt); PF01202 (SKI); PF01487 (DHquinase_I); PF01761 (DHQ_synthase); PF08501 (Shikimate_dh_N); PF01381 (HTH_3); PF01488 (Shikimate_DH); PF02153 (PDH); PF02224 (Cytidylate_kin); PF00501 (PF00501); PF13193 (PF13193); PF13560 (HTH_31); PF01817 (CM_2); PF00623 (RNA_pol_Rpb1_2); PF01885 (PF01885); PF03159 (XRN_N); PF04983 (RNA_pol_Rpb1_3); PF04997 (RNA_pol_Rpb1_1); PF04998 (RNA_pol_Rpb1_5); PF05000 (RNA_pol_Rpb1_4); PF13419 (HAD_2); PF00011 (HSP20); PF00132 (Hexapep); PF00443 (UCH); PF00616 (RasGAP); PF00670 (AdoHcyase_NAD); PF01019 (G_glu_transpept); PF01048 (PNP_UDP_1 and PF01134 (GIDA).

| Domains | Archaea | Bacteria | Eukaryota | Fungi | Viridiplantae | Stramenopiles |
|---|---|---|---|---|---|---|
| **EPSP_synthase** | **181** | **9052** | **623** | **412** | **133** | **26** |
| Chorismate_synt | 1 | 211 | 417 | 374 | 2 | 19 |
| SKI | 0 | 126 | 409 | 371 | 1 | 18 |
| DHquinase_I | 0 | 87 | 403 | 368 | 0 | 17 |
| DHQ_synthase | 0 | 17 | 398 | 368 | 0 | 15 |
| Shikimate_dh_N | 0 | 17 | 155 | 136 | 0 | 14 |
| HTH_3 | 0 | 9 | 2 | 2 | 0 | 0 |
| Shikimate_DH | 0 | 8 | 2 | 2 | 0 | 0 |
| PDH | 0 | 4 | 2 | 2 | 0 | 0 |
| Cytidylate_kin | 0 | 4 | 2 | 2 | 0 | 0 |

| | | | | | | |
|---|---|---|---|---|---|---|
| PF00501 | 0 | 3 | 2 | 2 | 0 | 0 |
| PF13193 | 0 | 3 | 2 | 2 | 0 | 0 |
| HTH_31 | 0 | 2 | 2 | 1 | 0 | 0 |
| CM_2 | 0 | 1 | 1 | 1 | 0 | 0 |
| RNA_pol_Rpb1_2 | 0 | 1 | 1 | 1 | 0 | 0 |
| PF01885 | 0 | 1 | 1 | 1 | 0 | 0 |
| XRN_N | 0 | 1 | 1 | 1 | 0 | 0 |
| RNA_pol_Rpb1_3 | 0 | 1 | 1 | 1 | 0 | 0 |
| RNA_pol_Rpb1_1 | 0 | 1 | 1 | 1 | 0 | 0 |
| RNA_pol_Rpb1_5 | 0 | 1 | 1 | 1 | 0 | 0 |
| RNA_pol_Rpb1_4 | 0 | 1 | 1 | 1 | 0 | 0 |
| HAD_2 | 0 | 0 | 1 | 1 | 0 | 0 |
| HSP20 | 0 | 0 | 1 | 0 | 0 | 0 |
| Hexapep | 0 | 0 | 1 | 0 | 0 | 0 |
| UCH | 0 | 0 | 1 | 0 | 0 | 0 |
| RasGAP | 0 | 0 | 1 | 0 | 0 | 0 |
| AdoHcyase_NAD | 0 | 0 | 1 | 0 | 0 | 0 |
| G_glu_transpept | 0 | 0 | 1 | 0 | 0 | 0 |
| PNP_UDP_1 | 0 | 0 | 1 | 0 | 0 | 0 |
| GIDA | 0 | 0 | 1 | 0 | 0 | 0 |

**Supplementary Table 6. Most frequent EPSPS associated domains in multidomain proteins.** FREQ is the frequency of the gene in samples, SP is the number of the gene in how many species they are found and FUNCTION in the description of product and if it is part of shikimate pathway.

| GENE | FREQ | SP | FUNCTION |
|---|---|---|---|
| **EPSPS** | **1448** | **8833** | **6$^{th}$ step of shikimate. pathway, produces EPSP Synthase** |
| SKI | 424 | 7716 | Start of pathway, produces phosphorylates shikimate |
| DHQ_synthase | 420 | 7429 | Second step of pathway, removes a phosphate from DHAP |

| | | | |
|---|---|---|---|
| DHquinase_I | 416 | 2073 | 3rd step of pathway, produces 3-dehydroquinate dehydratase |
| Shikimate_dh_N | 402 | 7658 | The substrate binding domain of shikimate dehydrogenase |
| HTH_3 | 218 | 9670 | Helix-turn-helix, a major structural motif capable of binding DNA |
| Shikimate_DH | 160 | 6719 | 4th step. Shikimate / quinate 5-dehydrogenase |
| PDH | 127 | 7136 | Prephenate dehydrogenases are part of Tyrosine biosynthesis. |
| Cytidylate_kin | 88 | 6874 | Kinase of cytidine 5'-monophosphate |
| PF13193 | 17 | 8031 | AMP-binding enzyme C-terminal domain for PF00501 |

**Supplementary table 7. List of common bacteria in the gut-microbiome (from supplementary tables 8, 5 and 12 provided by Qin et al. 2010) and their susceptibility to glyphosate.** These species or strains are specifically mentioned at Qin et al. (2010). N: Number of sequences; R: Sequences are resistant; S: Sequences are sensitive to glyphosate, U: sequences are unclassified and their sensitiveness is unknown.

| Species/strain | N | R / S | Class | Resistant Sequences (%) |
|---|---|---|---|---|
| ***Actinomyces odontolyticus* ATCC 17982** | 1 | S | Class I | 0% |
| ***Anaerofustis stercorihominis* DSM 17244** | 1 | S | Class I | 0% |
| ***Anaerotruncus colihominis*** | 5 | S | Class I | 0% |
| ***Bacteroides caccae*** | 5 | S | Class I | 0% |
| ***Bacteroides capillosus*** | 1 | S | Class I | 0% |
| ***Bacteroides cellulosilyticus*** | 4 | S | Class I | 0% |
| ***Bacteroides coprocola*** | 2 | S | Class I | 0% |
| ***Bacteroides coprophilus*** | 1 | S | Class I | 0% |
| ***Bacteroides coprophilus* DSM 18228** | 1 | S | Class I | 0% |
| ***Bacteroides dorei*** | 1 | S | Class I | 0% |
| ***Bacteroides dorei* DSM 17855** | 1 | S | Class I | 0% |
| ***Bacteroides eggerthii*** | 3 | S | Class I | 0% |
| ***Bacteroides finegoldii*** | 4 | S | Class I | 0% |
| ***Bacteroides fragilis* 3_1_12** | 1 | S | Class I | 0% |
| ***Bacteroides fragilis* NCTC 9343** | 1 | S | Class I | 0% |
| ***Bacteroides fragilis* YCH46** | 1 | S | Class I | 0% |
| ***Bacteroides intestinalis*** | 11 | S | Class I | 0% |
| ***Bacteroides ovatus*** | 14 | S | Class I | 0% |
| ***Bacteroides plebeius*** | 11 | S | Class I | 0% |
| ***Bacteroides sp.* 2_1_7** | 1 | S | Class I | 0% |
| ***Bacteroides sp.* 2_2_4** | 1 | S | Class I | 0% |

| | | | | |
|---|---|---|---|---|
| ***Bacteroides sp.* 3_2_5** | 1 | S | Class I | 0% |
| ***Bacteroides sp.* 4_3_47FAA** | 1 | S | Class I | 0% |
| ***Bacteroides sp.* 9_1_42FAA** | 2 | S | Class I | 0% |
| ***Bacteroides sp.* D1** | 1 | S | Class I | 0% |
| ***Bacteroides stercoris*** | 20 | S | Class I | 0% |
| ***Bacteroides uniformis*** | 10 | S | Class I | 0% |
| ***Bacteroides vulgatus* ATCC 8482** | 1 | S | Class I | 0% |
| ***Bifidobacterium adolescentis*** | 26 | 25×S 1×R | Class I; Class II | 4% |
| ***Bifidobacterium adolescentis* ATCC 15703** | 1 | S | Class I | 0% |
| ***Bifidobacterium adolescentis* L2-32** | 1 | S | Class I | 0% |
| ***Bifidobacterium bifidum* NCIMB 41171** | 1 | S | Class I | 0% |
| ***Bifidobacterium breve*** | 27 | S | Class I | 0% |
| ***Bifidobacterium breve* DSM 20213** | 1 | S | Class I | 0% |
| ***Bifidobacterium catenulatum*** | 7 | S | Class I | 0% |
| ***Bifidobacterium catenulatum* DSM 16992** | 1 | S | Class I | 0% |
| ***Bifidobacterium dentium*** | 5 | S | Class I | 0% |
| ***Bifidobacterium longum* DJO10A** | 1 | S | Class I | 0% |
| ***Bifidobacterium longum* NCC2705** | 1 | S | Class I | 0% |
| ***Bifidobacterium longum subsp. infantis* ATCC 15697** | 1 | S | Class I | 0% |
| ***Bifidobacterium longum subsp. infantis* ATCC 55813** | 1 | S | Class I | 0% |
| ***Bifidobacterium longum subsp. infantis* CCUG 52486** | 1 | S | Class I | 0% |
| ***Bifidobacterium pseudocatenulatum*** | 46 | 45×S<br>1×U | Class I<br>U | 0% |
| ***Catenibacterium mitsuokai*** | 3 | S | Class I | 0% |
| ***Citrobacter sp.* 30_2** | 1 | S | Class I | 0% |
| ***Citrobacter sp.* 30_3** | 1 | S | Class I | 0% |
| ***Citrobacter sp.* 30_4** | 1 | S | Class I | 0% |
| ***Citrobacter sp.* 30_5** | 1 | S | Class I | 0% |
| ***Citrobacter sp.* 30_6** | 1 | S | Class I | 0% |
| ***Clostridium leptum*** | 4 | S | Class I | 0% |
| ***Clostridium leptum* DSM 753** | 1 | U | U | 0% |
| ***Clostridium methylpentosum*** | 1 | S | Class I | 0% |
| ***Clostridium nexile*** | 1 | S | Class I | 0% |
| ***Collinsella aerofaciens*** | 9 | S | Class I | 0% |
| ***Desulfovibrio piger* ATCC 29098** | 1 | S | Class I | 0% |
| ***Enterobacter cancerogenus* ATCC 35316** | 1 | S | Class I | 0% |
| ***Escherichia coli* 536** | 1 | S | Class I | 0% |

| | | | | |
|---|---|---|---|---|
| ***Escherichia coli* APEC O1** | 1 | S | Class I | 0% |
| ***Escherichia coli* ATCC 8739** | 1 | S | Class I | 0% |
| ***Escherichia coli* CFT073** | 1 | S | Class I | 0% |
| ***Escherichia coli* ED1a** | 1 | S | Class I | 0% |
| ***Escherichia coli* HS** | 1 | S | Class I | 0% |
| ***Escherichia coli* IAI39** | 1 | S | Class I | 0% |
| ***Escherichia coli* O127:H6 str. E2348/69** | 1 | S | Class I | 0% |
| ***Escherichia coli* O157:H7 str. EC4115** | 1 | S | Class I | 0% |
| ***Escherichia coli* S88** | 1 | S | Class I | 0% |
| ***Escherichia coli* SE11** | 1 | S | Class I | 0% |
| ***Escherichia coli* SMS-3-5** | 1 | S | Class I | 0% |
| ***Escherichia coli* UTI89** | 1 | S | Class I | 0% |
| ***Eubacterium siraeum*** | 3 | S | Class I | 0% |
| ***Eubacterium siraeum* 70/3** | 1 | S | Class I | 0% |
| ***Faecalibacterium prausnitzii* M21/2** | 1 | S | Class I | 0% |
| ***Faecalibacterium prausnitzii* SL3 3** | 1 | S | Class I | 0% |
| ***Fusobacterium mortiferum* ATCC 9817** | 1 | S | Class I | 0% |
| ***Fusobacterium sp.* 4_1_13** | 1 | S | Class I | 0% |
| ***Fusobacterium ulcerans* ATCC 49185** | 1 | S | Class I | 0% |
| ***Fusobacterium varium* ATCC 27725** | 1 | S | Class I | 0% |
| ***Holdemania filiformis*** | 1 | S | Class I | 0% |
| ***Methanobrevibacter smithii*** | 6 | S | Class I | 0% |
| ***Methanobrevibacter smithii* ATCC 35061** | 1 | S | Class I | 0% |
| ***Mollicutes bacterium* D7** | 1 | S | Class I | 0% |
| ***Parabacteroides distasonis* ATCC 8503** | 1 | S | Class I | 0% |
| ***Parabacteroides johnsonii*** | 4 | S | Class I | 0% |
| ***Parabacteroides merdae*** | 8 | S | Class I | 0% |
| ***Prevotella copri*** | 27 | S | Class I | 0% |
| ***Providencia alcalifaciens* DSM 30120** | 1 | S | Class I | 0% |
| ***Providencia rustigianii* DSM 4541** | 1 | S | Class I | 0% |
| ***Providencia stuartii* ATCC 25827** | 1 | S | Class I | 0% |
| ***Ruminococcus bromii*** | 13 | S | Class I | 0% |
| ***Ruminococcus bromii* L2-63** | 1 | S | Class I | 0% |
| ***Subdoligranulum variabile*** | 2 | S | Class I | 0% |
| ***Subdoligranulum variabile* DSM 15176** | 1 | S | Class I | 0% |
| ***Alistipes putredinis*** | 3 | R | Class III | 100% |
| ***Anaerobaculum hydrogeniformans* ATCC BAA-1850** | 1 | R | Class II | 100% |
| ***Blautia hansenii*** | 3 | R | Class II | 100% |

| | | | | |
|---|---|---|---|---|
| ***Bryantella formatexigens*** | 1 | R | Class II | 100% |
| ***Clostridium asparagiforme*** | 1 | R | Class II | 100% |
| ***Clostridium asparagiforme* DSM 15981** | 1 | R | Class II | 100% |
| ***Clostridium bartlettii*** | 2 | R | Class II | 100% |
| ***Clostridium scindens*** | 2 | R | Class II | 100% |
| ***Clostridium sp* L2-50** | 2 | 1×R<br>1×U | Class II<br>U | 50% |
| ***Clostridium sp* SS2-1** | 2 | R | Class II | 100% |
| ***Clostridium symbiosum* ATCC 14940** | 1 | R | Class II | 100% |
| ***Coprococcus comes*** | 8 | R | Class II | 100% |
| ***Coprococcus comes* ATCC 27758** | 1 | R | Class II | 100% |
| ***Coprococcus eutactus*** | 6 | R | Class II | 100% |
| ***Dorea formicigenerans*** | 18 | R | Class II | 100% |
| ***Dorea longicatena*** | 11 | R | Class II | 100% |
| ***Enterococcus casseliflavus* EC10** | 1 | R | Class II | 100% |
| ***Enterococcus casseliflavus* EC20** | 1 | R | Class II | 100% |
| ***Eubacterium hallii/Anaerobutycum hallii*** | 9 | R | Class II | 100% |
| ***Helicobacter canadensis* MIT 98-5491** | 1 | R | Class II | 100% |
| ***Helicobacter cinaedi* CCUG 18818** | 2 | R | Class II | 100% |
| ***Helicobacter pullorum* MIT 98-5489** | 1 | R | Class II | 100% |
| ***Helicobacter winghamensis* ATCC BAA-430** | 1 | R | Class II | 100% |
| ***Lactobacillus brevis subsp. gravesensis* ATCC 27305** | 1 | R | Class II | 100% |
| ***Lactobacillus buchneri* ATCC 11577** | 1 | R | Class II | 100% |
| ***Lactobacillus hilgardii* ATCC 8290** | 1 | R | Class II | 100% |
| ***Lactobacillus plantarum subsp. plantarum* ATCC 14917** | 1 | R | Class II | 100% |
| ***Lactobacillus plantarum* WCFS1** | 1 | R | Class II | 100% |
| ***Lactobacillus ruminis* ATCC 25644** | 2 | R | Class II | 100% |
| ***Listeria grayi* DSM 20601** | 1 | R | Class II | 100% |
| ***Mitsuokella multacida*** | 2 | R | Class II | 100% |
| ***Mitsuokella multacida* DSM 20544** | 1 | R | Class II | 100% |
| ***Ruminococcus gnavus*** | 16 | R | Class II | 100% |
| ***Ruminococcus gnavus* ATCC 29149** | 1 | R | Class II | 100% |
| ***Ruminococcus lactaris*** | 2 | R | Class II | 100% |
| ***Ruminococcus lactaris* ATCC 29176** | 1 | R | Class II | 100% |
| ***Ruminococcus obeum*** | 17 | R | Class II | 100% |
| ***Ruminococcus obeum* A2-162** | 1 | R | Class II | 100% |
| ***Ruminococcus torques*** | 7 | 6×R<br>1×U | Class II<br>U | 86% |

| | | | | |
|---|---|---|---|---|
| ***Ruminococcus torques* L2-14** | 1 | R | Class II | 100% |
| ***Bacteroides pectinophilus* ATCC 43243** | 1 | U | U | 0% |
| ***Butyrivibrio crossotus*** | 2 | U | U | 0% |
| ***Clostridium spiroforme* DSM 1552** | 1 | U | U | 0% |
| ***Eubacterium rectale* DSM 17629** | 1 | U | U | 0% |
| ***Eubacterium rectale* M104 1** | 1 | U | U | 0% |
| ***Eubacterium ventriosum*** | 7 | U | U | 0% |
| ***Fusobacterium sp.* 2_1_31** | 1 | U | U | 0% |
| ***Fusobacterium sp.* D11** | 1 | U | U | 0% |
| ***Fusobacterium sp.* D12** | 1 | U | U | 0% |
| ***Proteus penneri* ATCC 35198** | 1 | U | U | 0% |
| ***Providencia rettgeri* DSM 1131** | 1 | U | U | 0% |
| ***Roseburia intestinalis* L1-82** | 1 | U | U | 0% |
| ***Roseburia intestinalis* M50 1** | 1 | U | U | 0% |

#### SUPPLEMENTARY FIGURES

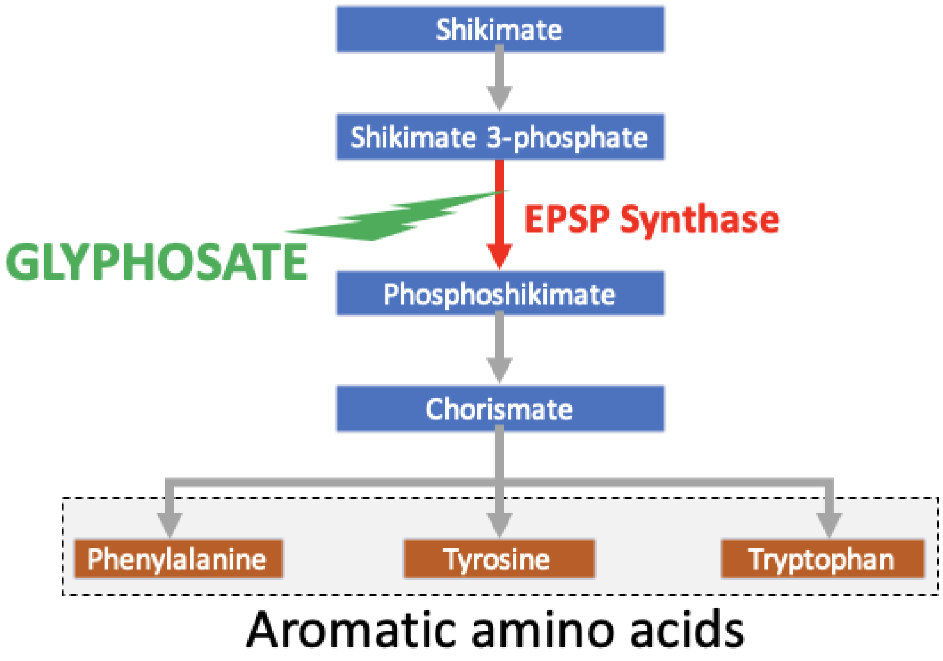


**Supplementary figure 1. Scheme of the effect of glyphosate on its target enzyme 5-enolpyruvylshikimate-3-phosphate synthase (EPSPS).** Glyphosate inhibits the key enzyme of the shikimate pathway and blocks the production of three essential amino acids (phenylalanine, tyrosine and tryptophan).

```
Colouring of similarity:
BAD AVG GOOD

                                  Colouring of active site positions:

vcEPSPS Class I   :  87           class Ia, class Ib, both
cbEPSPS Class II  :  85           class II
bvEPSPS Class III :  86           class III, the 18 motifs are squared
sdEPSPS Class IV  :  86           class IV
cons              :  87

vcEPSPS Class I   MES----LTLQ-PIELISGEVNLPGSKSVSNRALLLAALASGTTRLTNLLDSDDIRHMLNALT
cbEPSPS Class II  MDY----QT-I-PSQGLSGEICVPGDKSISHRAVLLAAIAEGQTQVDGFLMGADNLAMVSALQ
bvEPSPS Class III MMMGRAKLTIIPPGKPLTGRAMPPGSKSITNRALLLAGLAKGTSRLTGALKSDDTRYMAEALR
sdEPSPS Class IV  MP----------------VADIPGSKSITARALFLAAADGVTTLVRPLRSDDTEGFAEGLA
cons              *                      **.**:: **::**. *.* : :    * . *    : ..*

vcEPSPS Class I   KLGVNYRLSADKTTCEVEGLGQAFHTTQPLELFLGNAGTAMRPLAAALCLGQGDYVLTGEPRM
cbEPSPS Class II  QMGASIQVIEDENILVVEGVGMTGLQAPPEALDCGNSGTAIRLLSGLLAGQPFNTVLTGDSSL
bvEPSPS Class III AMGVTIDEPD-DTTFIVKGSGKLQP--PAAPLFLGNAGTATRFLTAAAALVDGKVIVDGDAHM
sdEPSPS Class IV  RLGYRVGRTP--DSWQVDGRPQGPA-VAEADVYCRDGATTARFLPTLAAAGHGTYRFDASEQM
cons               :*               *.*          :    :..*: * *.    .           .  ..  :

vcEPSPS Class I   KERPIGHLVDALRQAGAQIEYLEQENFPPLRIQGTGLQAG-TVTIDGSISSQFLTAFLMSAPL
cbEPSPS Class II  QRRPMKRIIDPLTLMGAKIDS--TGNVPPLKIYGNPRLTGIHYQL-PMASAQVKSCLLLAGLY
bvEPSPS Class III RKRPIGPLVDALRSLGIDASA--ETGCPPVTINGTGRFEASRVQIDGGLSSQYVSALLMMAAG
sdEPSPS Class IV  RRRPLLPLTRALRELGVDLRHEERDGHHPLTVRAAGV-AGGEVTLDAGQSSQYLTALLLLGPL
cons              :.**:  :  .*   * .     .  *: : .         .      :      *:*  :.:*: .

vcEPSPS Class I   AQGKVTIKIVGEL-VSKPYIDITLHIMEQFGVQVINHDY-QEFVIPAGQSYVSPGQFLVEGDA
cbEPSPS Class II  ARGKTCIT---E---PAPSRDHTERLLKHFHYTLQ-KDK-QSICVSGGGKL-KANDISIPGDI
bvEPSPS Class III GDRAVDVELLGEHIGALGYIDLTVAAMRAFGAKVE-RVSPVAWRVEPTG-Y-HAADFVIEPDA
sdEPSPS Class IV  TEKGLRIHV-TDL-VSVPYIEITLAMMRAFGVEVT-REG-HDFVVPPGG-Y-RATTYAIEPDA
cons                    :    :   .     : *    :. *    :  :         :       .    :  *

vcEPSPS Class I   SSASYFLAAAAIKGG-EVKVTGIGKNSIQGDIQFADALEKMGAQIEWGD---------DYVIA
cbEPSPS Class II  SSAAFFIVAATITPGSAIRLCRVGVNPT--RLGVINLLKMMGADIEVTHYTEKNEEPTADITV
bvEPSPS Class III SAATYLWAAEVLSGG-KIDLGTPAEQFSQPDAKAYDLISKFP---------------------
sdEPSPS Class IV  STSSYFFAAAALSGG-EVTVPGLGEGALQGDLGFVDVLRRMGAEVEIGA---------DRTTV
cons              *::::: .* .:. *  : :    .            : :  :

vcEPSPS Class I   R-RGELNAVDLDFNHIP---DAAMTIATTALFAKGTTAIRNVYNWRVKETDRLAAMATELRKV
cbEPSPS Class II  R-HARLKGIDIPPDQVPLTIDEFPVLLIAAAVAQGKTVLRDAAELRVKETDRIAAMVDGLQKL
bvEPSPS Class III ----HLPAV-IDGSQMQ---DAIPTLAVLAAFNEMPVRFVGIENLRVKECDRIRALSSGLSRI
sdEPSPS Class IV  RGTGELRGLTVNMRDIS---DTMPTLAAIAPFASGPVRIEDVANTRVKECDRLEACAENLRRL
cons                  .* .: :    .:     *    .:     *  .  .    .  : .   : **** **: *      * ::

vcEPSPS Class I   G--ATVEEGEDFIVITPPT---KLIHAAIDTYDDHRMAMCFSLVA-LSDTPVTINDPKCTSKT
cbEPSPS Class II  G--IAAESLPDGVIIQGG----TLEGGEVNSYDDHRIAMAFAVAGTLAKGPVRIRNCDNVKTS
bvEPSPS Class III VPNLGTEEGDDLIIASDPSLAGKILTAEIDSFADHRIAMSFALAG-LKIGGITILDPDCVAKT
sdEPSPS Class IV  G--VRVETGPDWIEIHPGA---TPTGAEIKTYGDHRIVMSFAVTG-LRVPGISFDDPGCVRKT
cons                 .*   * :        .    .  :.::  ***:.*.*::.. *      :  :  :     . .:

vcEPSPS Class I   FPDYFDKFAQLSR-----------
cbEPSPS Class II  FPNFVELANEVGMNVKGVRGRGGF
bvEPSPS Class III FPSYWNVLSSLGVAYED-------
sdEPSPS Class IV  FPGFHEEFGALRAR--L-------
cons              **.: :    :
```

**Supplementary figure 2.** Multiple sequence alignment made with T-Coffee [3,4] of the four EPSPS reference sequences.

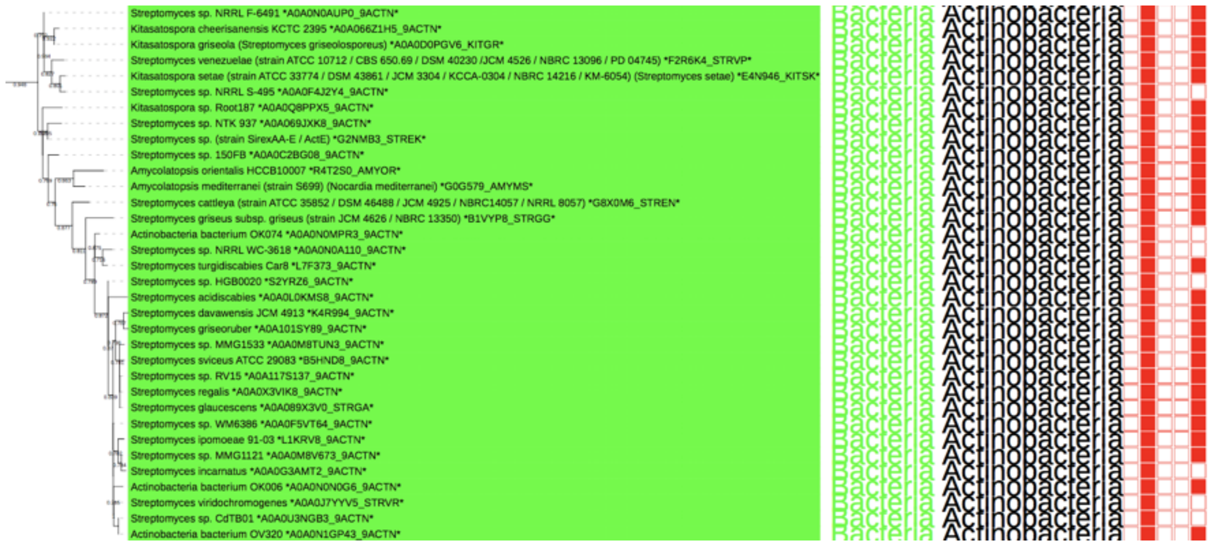


**Supplementary figure 3. All EPSPS class IV belong to one single clade of actinobacteria species.**

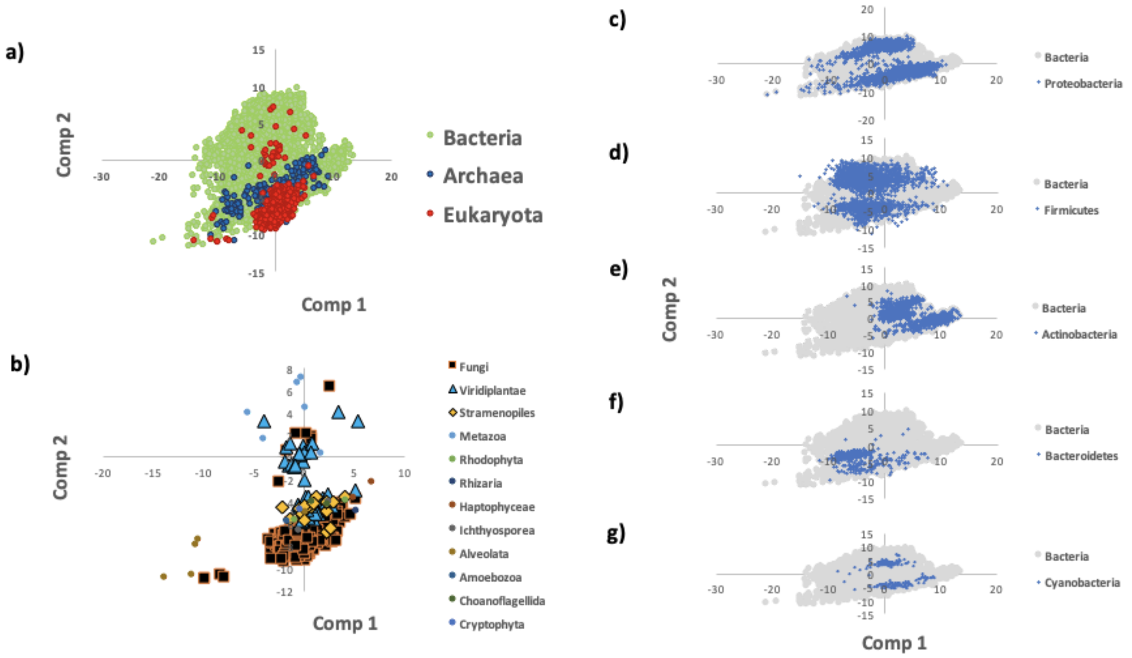


**Supplementary figure 4. Principal Components Analysis (PCA) plot of dipeptides by taxonomy**
First and second component of the PCA of ~10,000 EPSPS proteins
a) Archaea (blue), bacteria (green) and eukaryotes (red)
b) Eukaryotes c-g) Prokaryotes

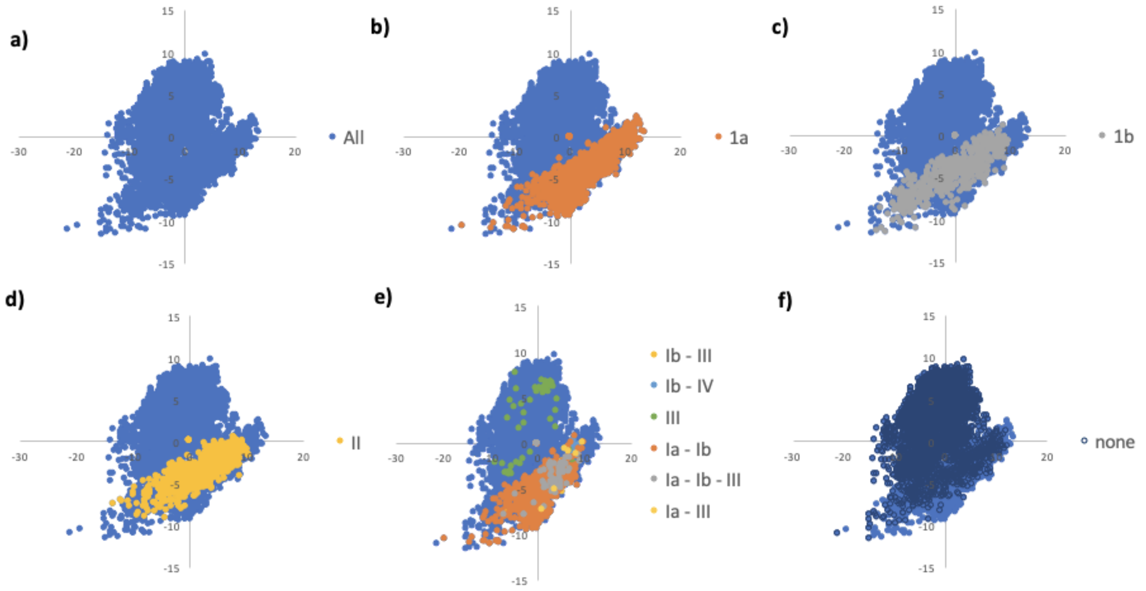


**Supplementary figure 5. Principal Components Analysis (PCA) plot of dipeptides by EPSPS class**
First and second component of the PCA of ~10,000 EPSPS proteins
a) All
b) Class I alpha
c) Class I beta
d) Class II
e) Multiple classes
f) Unclassified

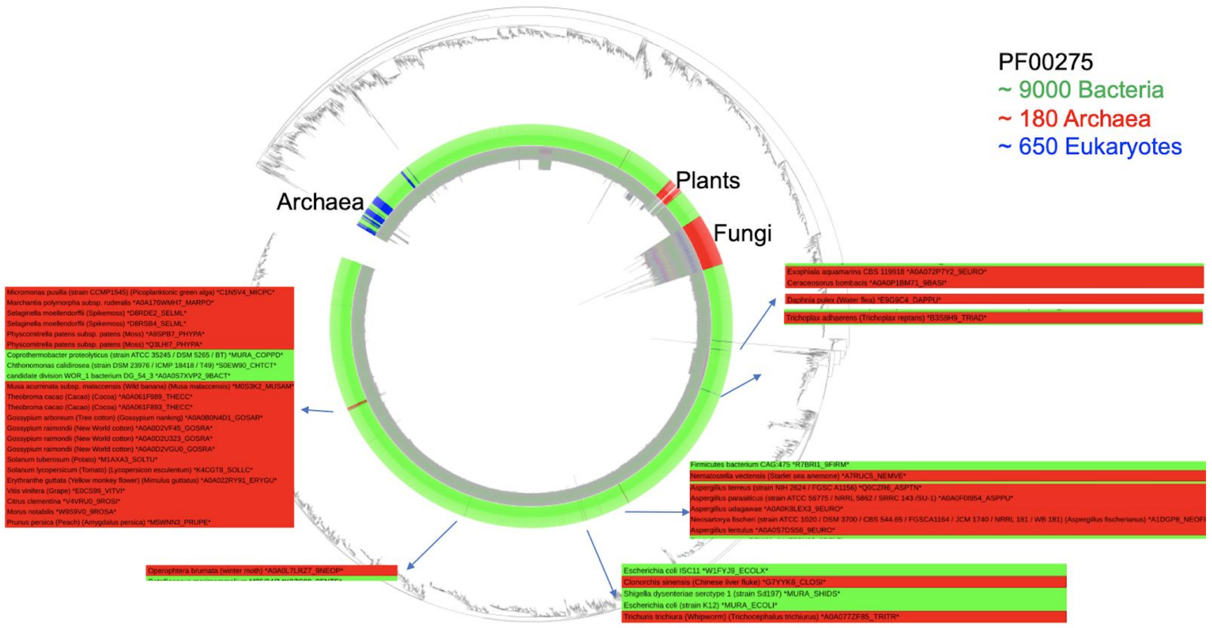


**Supplementary figure 6. Eukaryotic species across the species tree (Figure 1a).**

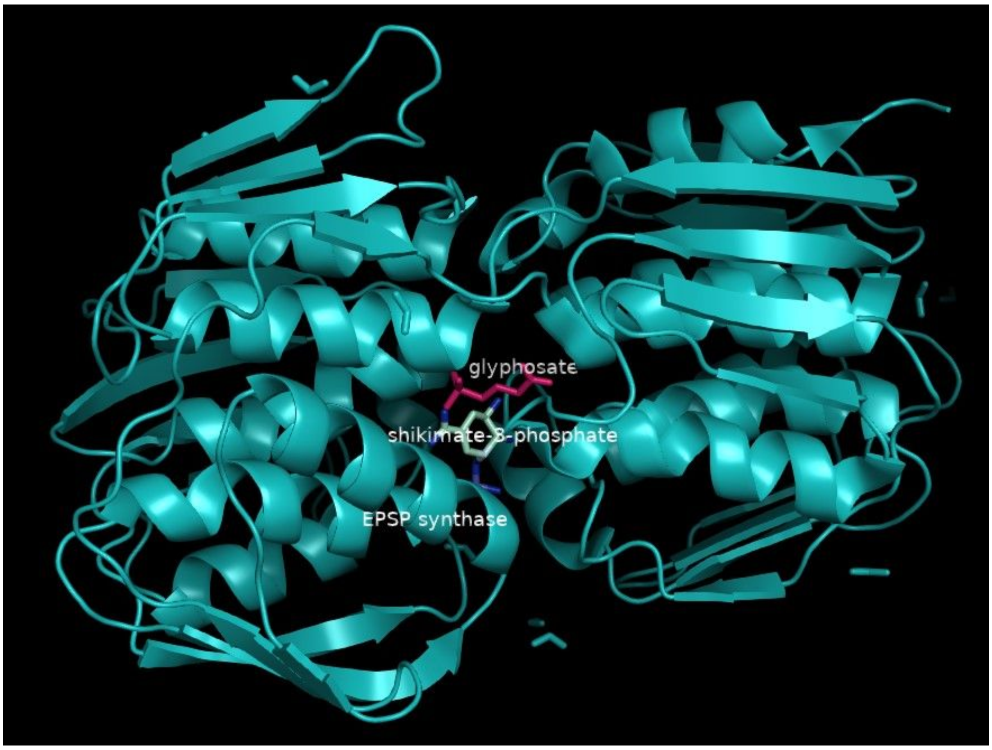


**Supplementary figure 7.** A 3D-structure illustration of the EPSP synthase of *E. coli* (RCSB PDB code: 1G6S) liganded with shikimate-3-phosphate and glyphosate (The PyMOL Molecular Graphics System, Version 1.2r3pre, Schrödinger, LLC).
